# Immunity-and-Matrix-Regulatory Cells Derived from Human Embryonic Stem Cells Safely and Effectively Treat Mouse Lung Injury and Fibrosis

**DOI:** 10.1101/2020.04.15.042119

**Authors:** Jun Wu, Dingyun Song, Zhongwen Li, Baojie Guo, Yani Xiao, Wenjing Liu, Lingmin Liang, Chunjing Feng, Tingting Gao, Yanxia Chen, Ying Li, Zai Wang, Jianyan Wen, Shengnan Yang, Peipei Liu, Lei Wang, Yukai Wang, Liang Peng, Glyn Nigel Stacey, Zheng Hu, Guihai Feng, Wei Li, Yan Huo, Ronghua Jin, Ng Shyh-Chang, Qi Zhou, Liu Wang, Baoyang Hu, Huaping Dai, Jie Hao

## Abstract

Lung injury and fibrosis represent the most significant outcomes of severe and acute lung disorders, including COVID-19. However, there are still no effective drugs to treat lung injury and fibrosis. In this study, we report the generation of clinical-grade human embryonic stem cells (hESCs)-derived immunity- and matrix-regulatory cells (IMRCs) produced under good manufacturing practice (GMP) requirements, that can treat lung injury and fibrosis in vivo. We generate IMRCs by sequentially differentiating hESCs with serum-free reagents. IMRCs possess a unique gene expression profile distinct from umbilical cord mesenchymal stem cells (UCMSCs), such as higher levels of proliferative, immunomodulatory and anti-fibrotic genes. Moreover, intravenous delivery of IMRCs inhibits both pulmonary inflammation and fibrosis in mouse models of lung injury, and significantly improves the survival rate of the recipient mice in a dose-dependent manner, likely through paracrine regulatory mechanisms. IMRCs are superior to both primary UCMSCs and FDA-approved pirfenidone, with an excellent efficacy and safety profile in mice and monkeys. In light of public health crises involving pneumonia, acute lung injury (ALI) and acute respiratory distress syndrome (ARDS), our findings suggest that IMRCs are ready for clinical trials on lung disorders.

## Introduction

Lung injury is a serious threat to human health. It is caused by many chemical, physical and biological agents, resulting in pneumonia, lung inflammation, and fibrosis. In a comprehensive analysis, the World Health Organization ranks lung diseases as the second largest category, and predicts that by 2020, about one fifth of human deaths will be attributed to lung diseases^1^. Acute lung injury (ALI) is the injury of alveolar epithelial cells and capillary endothelial cells caused by various direct and indirect injury factors, resulting in diffuse pulmonary interstitial and alveolar edema, and acute hypoxic respiratory insufficiency. Pathophysiologically, its characteristics include decreased lung volume, decreased lung extensibility and imbalance of the ventilation/blood flow ratio. When ALI develops into the severe stage (oxygenation index < 200), it is called acute respiratory distress syndrome (ARDS). Globally, there are more than 3 million patients with ARDS annually, accounting for 10% of intensive care unit (ICU) admissions^2^. ALI/ARDS patients have an average mortality rate of 35-46%^3,4^. In late ARDS, pulmonary fibrosis (PF) develops due to persistent alveolar injury, repeated destruction, repair, reconstruction and over-deposition of extracellular matrix (ECM)^5^, leading to progressive lung scars and common interstitial pneumonia. In general, idiopathic pulmonary fibrosis (IPF) patients show very poor prognosis, with a median survival time of 3-5 years after diagnosis^6^. It has also been reported that the degree of pulmonary fibrosis is closely related to the mortality of ARDS^7,8^. Importantly, there are still no effective FDA-approved drugs to treat ALI, ARDS or IPF hitherto, and most experimental drugs are still in the midst of Phase II or Phase III clinical studies^2, 9,10^.

Stem cell therapy is an emerging treatment modality being used to cure various inflammatory and/or degenerative diseases, including ALI and PF. In particular, mesenchymal stem cells (MSCs) have been tested as an intravenous infusion therapy for ALI and PF^11,12^. Preclinical and clinical studies of pulmonary fibrosis have shown that, after infusion, MSCs could migrate to the sites of lung injury, inhibit inflammation and improve recovery^13-15^. However, clinical studies with primary MSCs derived from the umbilical cord, bone marrow or adipose tissue have been hampered by the lack of available donors, limited cell numbers from each donor, donor and tissue heterogeneity, inconsistent cell quality and the lack of standardized cell preparations. Moreover, primary MSCs show limited self-renewal capacities and finite lifespans. MSC-like cells derived from human embryonic stem (hESCs)^16^ could solve these problems^17-19^, but the use of ill-defined and variable components of animal origin, such as fetal bovine serum (FBS) or OP9 feeder cells, may greatly compromise their consistency, safety and clinical applicability. Furthermore, these hESC-derived MSC-like cells have not been deeply characterized for their immunomodulatory functions in disease models.

In this study, we demonstrate that our hESC-derived MSC-like cell population have unique abilities in modulating the immunity and regulating extracellular matrix production, compared to regular MSC populations. Therefore, we named it as immunity- and matrix-regulatory cells (IMRCs). One aspect of IMRCs is that they resemble MSCs in their capacity for self-renewal and tri-lineage differentiation. However, compared to primary umbilical cord mesenchymal stem cells (UCMSCs), IMRCs also displayed a higher consistency in quality, stronger immunomodulatory and anti-fibrotic functions, and a robust ability to treat lung injury and fibrosis in vivo.

## Materials and methods

### Generation of hESC-derived IMRCs

The hESC-derived IMRCs were generated by passaging cells that are migrating out from human embryoid bodies (hEBs), with serum-free reagents. The clinical hESC line (Q-CTS-hESC-2) was prepared as described previously^20^. Clinical hESCs were maintained in commercially available Essential 8™ (E8) basal medium (Gibco, Carlsbad, MO, USA; A15169-01) with Essential 8™ Supplement (Gibco, A15171-01) on vitronectin-coated plates (1 µg/cm^2^). To generate embryoid bodies (hEBs), hESCs were dissociated into small clumps by incubating at 37 °C for 5 min with 1 mg/mL dispase (Gibco, 17105-04) and cultured to form hEBs for 5 days in KO-DMEM (Gibco, A12861-01) supplemented with 20% KOSR (Gibco, A3020902), 1 × L-glutamine (Gibco, A12860-01), 1 × NEAA (Gibco, 11140050) and 10 ng/mL bFGF (Thermo, 13256029). Then, hEBs were transferred onto vitronectin-coated plates and cultured for 14 additional days. During this period, the outgrowth from the hEBs occurred. The outgrowth cells were dissociated by using Tryple (Gibco, A12859-01) and passaged at a low cell density of 1 × 10^4^ cells per cm^2^ in “IMRCs Medium” consisting of α-MEM (Gibco, 12561-049) supplemented with 5% KOSR, 1% Ultroser G (Pall corporation, New York, NY, USA; 15950-017), 1 × L-glutamine, 1 × NEAA, 5 ng/mL bFGF and 5 ng/mL TGF-β (Peprotech, 96-100-21-10). Cells were passaged upon reaching approximately 80% confluence in IMRCs Medium. After 5 passages in culture, differentiated cultures were harvested for characterization. By this time, IMRCs displayed a fibroblastic morphology, expressed canonical “MSC-specific” surface markers including CD73, CD90, CD105 and CD29 and were negative for typical hematopoietic markers (CD45, CD34, HLA-DR)^21,22^.

### Isolation and culture of UCMSCs

Healthy full-term human placental samples were collected according to the policy of the Ethics Committee of the 306^th^ Hospital of the Chinese People’s Liberation Army, Beijing, China. Written informed consents were obtained from all donors before this study. All the samples were used in accordance with standard experimental protocols approved by the Animal and Medical Ethical Committee of Institute of Zoology, Chinese Academy of Sciences, Beijing, China.

Briefly, the newborn umbilical cords of full-term pregnancies were obtained from the clinic and washed by PBS to remove all remnant blood. Then, the umbilical cords were cut to approximately 2 mm in size after removing the artery and vein. The pieces of umbilical cords were directly transferred into a 10 cm^2^ culture flasks in α-MEM supplemented with 5% KOSR, 1% Ultroser G, 1 × L-glutamine, 1 × NEAA and 5 ng/mL bFGF, and cultured in an atmosphere of 5% CO_2_ at 37 °C. UCMSCs were passaged upon reaching approximately 80% confluence. After 5 passages in culture, cells were harvested for characterization.

### Cell culture

The clinical hESC line (Q-CTS-hESC-2) was prepared as described previously^20^. Clinical hESCs were maintained in the commercially available Essential 8™ (E8) basal medium on vitronectin-coated plates. Cells were passaged every 5 or 6 days using Versene (Gibco, A4239101). Clinical hESCs were tested weekly for mycoplasma contamination using a Myco-detection Kit (InvivoGen, San Diego, CA, USA; rep-pt1) and endotoxin contamination was tested using a ToxinSensor™ Chromogenic LAL Endotoxin Assay Kit (GenScript, Piscataway, NJ, USA; L00350). All cultures were maintained at 37 °C, 5% CO_2_ and atmospheric O_2_ in a humidified incubator (Thermo Fisher Scientific, Waltham, MA, USA). A549 cells were purchased from Beina Biology (Shanghai, China). The cells were cultured in Ham’s F-12K medium (Gibco, 21127022) with 10% fetal bovine serum (FBS; Gibco, 10091-148) at 37 °C in 5% CO_2_, and were treated with 10 ng/mL TGF-β1 (R&D systems, 240-B) or IMRC conditioned medium where specified. To prepare the conditioned medium of IMRCs, IMRCs were cultured to 50% confluence in culture dishes, then, were changed to new medium. The conditioned medium was harvested after incubating IMRCs for 24 h.

### Karyotyping

At passage 5, IMRCs were harvested. Karyotype analysis and G-binding were conducted at the Chinese Academy of Medical Science & Peking Union Medical College (Beijing, China).

### RNA-seq library preparation and data analysis

Total RNA was isolated from hESCs, IMRCs and UCMSCs by Trizol (Invitrogen, Waltham, MA, USA; 15596018). RNA-seq libraries were prepared with the NEBNext^®^Ultra™ RNA Library Prep Kit for Illumina^®^. Sequencing was performed on an Illumina HiSeq X-Ten sequencer with 150 bp paired-end sequencing reaction. The bulk RNA-Seq data for hESCs were downloaded from GEO database^23^. All RNA-sequencing data analysis was performed with Hisat2 (version 2.1.0)^24^ and Cufflinks (version 2.2.1)^25^ using the UCSC hg19 annotation with default settings. Reads with unique genome locations and genes with no less than 1 FPKM in at least one sample were used for the next step of analysis. Two-fold change was used as the threshold to filter for differentially expressed genes (DEGs) i.e. increased or decreased expression of less than 2-fold was not considered significant. The Gene Ontology analysis for differentially expressed genes was performed by DAVID (version 6.8)^26^. Clustering, heatmap, Venn diagram and scatterplots analysis were performed with hierarchical cluster and heatmap.2 functions in R. Pearson correlation coefficient was calculated by cor.test in R. Gene set enrichment analysis were performed by GSEA^27^.

### Flow cytometry

Cells were harvested and blocked with 2% bovine serum albumin (BSA; Sigma-Aldrich, B2064) for 20 min at room temperature. Then, the cells were stained with fluorescein-conjugated antibodies for 40 min at room temperature in 1% BSA. After incubation, cells were washed 3 times and analyzed with MoFlo (Beckman, Brea, CA, USA) and associated software. The antibodies used for flow cytometry were as follows: PE-conjugated mouse anti-human CD29 (Biolegend, San Diego, CA, USA; 303004), PE-conjugated mouse anti-human CD73 (BD Biosciences, San Jose, CA, USA; 550257), PE-conjugated mouse anti-human CD90 (eBioscience, San Diego, CA, USA; 12-0909-42), PE-conjugated mouse anti-human CD105 (Biolegend, 323206), PE-conjugated mouse anti-human CD34 (BD Biosciences, 555822), PE-conjugated mouse anti-human CD45 (BD Biosciences, 560957), PE-conjugated mouse anti-human HLA-DR (BD Biosciences, 555561), and the PE-conjugated mouse IgG1 (BD Biosciences, 551436) as an isotype control.

### Trilineage differentiation

Osteogenic, adipogenic, and chondrogenic differentiation experiments were performed following the instructions of the human mesenchymal stem cell functional identification kit (R&D systems, SC006). For osteogenic differentiation, 4.2 × 10^3^ cells were seeded per well in 96-well plates. When cells reached 50-70% confluency, the medium was replaced with osteogenic differentiation medium and kept for 3 weeks. To assess osteogenic differentiation, immunofluorescence (Osteocalcin) and Alizarin Red S (Sigma-Aldrich, A5533) staining was performed for the calcium-rich extracellular matrix. For adipogenic differentiation, cells were seeded into a 24-well plate at the density of 3.7 × 10^4^ cells/well, and maintained in culture medium until 100% confluency. Then, cells were cultured in adipogenic differentiation medium for 3 weeks. Lipid droplets of the resultant differentiated cells were detected using immunofluorescence (FABP-4) and Oil red O (Sigma-Aldrich, O0625) staining. For chondrogenic differentiation, 2.5 × 10^5^ cells resuspended in chondrogenic differentiation medium were centrifuged for 5 min at 200 g in a 15-mL conical tube (Corning, NY, USA). Then, cells were cultured for 3 weeks. After 3 weeks, chondrogenic pellet was harvested and fixed in 4% paraformaldehyde (Aladdin Chemical Co., Ltd., Shanghai, China; C104190). Cryosectioning was performed by OTF Cryostat (Leica, Germany) and 10 µm sections were stained with immunofluorescence (Aggrecan) and Alcian Blue (Sigma-Aldrich, A5268) staining.

### Growth curve

IMRCs and UCMSCs were seeded into 96-well plates at a density of 8 × 10^3^ cells per well. Cell Counting Kit-8 (CCK8; Thermo Fisher Scientific, CK04) was used to measure the cells’ growth. CCK8 solution was added at day 1, day 3, day 5 and day 7 by following the CCK8 kit manufacturer’s protocol. After 2 h of incubation, absorbance was measured at 450 nm.

### Cell diameter analysis

The IMRCs and UCMSCs were harvested at passage 5 and the cell diameters were measured with Image J software (NIH) software analysis. Normal Distribution curves were fitted to the cell diameter data to analyze the cell size distribution characteristics.

### Cell viability test

The IMRCs and UCMSCs were cryopreserved in a published clinical injection^28^ for 15 days in liquid nitrogen. After complete recovery of the cells, cells were suspended in clinical injection and remained under 4 °C. Then, cell viability was determined using 0.4% (w/v) Trypan Blue (Gibco, 15250061) at different time points by a Cell Counter (Invitrogen, Countess II) using the average of 3 replicates.

### Single-cell RNA-seq library preparation & sequencing

IMRCs and UCMSCs were harvested and suspended in PBS. Then, cell suspensions were loaded onto the Chromium Single Cell Controller (10 × Genomics) to generate single Gel Beads-in-Emulsion (GEMs) by using Single Cell 30 Library and Gel Bead Kit V2 (10 × Genomics, 120237). Cells were lysed and the released RNA was barcoded through reverse transcription in individual GEMs. Following reverse transcription, cDNAs with both barcodes were amplified, and a library was constructed using the Single Cell 30 Reagent Kit (v2 chemistry) for each sample following the manufacture’s introduction (performed by CapitalBio Technology, Beijing). Sequencing was performed on an Illumina NovaSeq 6000 System in the 2 × 150 bp paired-end mode. Raw files were processed with Cell Ranger using default mapping arguments. Gene expression analysis and cell type identification were performed by Seurat V3.1. Briefly, after normalizing and quality control, the tSNE was used for non-linear dimensional reduction. The figures were produced by DimPlot and VlnPlot functions in Seurat.

### Cell culture with IFN-γ treatment

Cells at passage 4 were digested and seeded into 6-well plates. IMRCs were seeded at a density of 3 × 10^5^ cells/well, and UCMSCs were seeded at a density of 2 × 10^5^ cells/well. After adherence for 24 h, IFN-γ (0, 25, 50, 100 ng/mL) (R&D systems, 285-IF) was added into the culture medium to culture for 24 h. Cells and conditioned medium were harvested after IFN-γ stimulation for qPCR and cytokine analysis.

### Cytokine analysis

Conditioned medium of UCMSCs and IMRCs were collected after IFN-γ (100 ng/mL) stimulation for 24 h. Forty-eight cytokines including FGF basic, Eotaxin (also known as CCL11), G-CSF (CSF3), GMCSF (CSF2), IFN-γ, IL-1β, IL-1RA, IL-2, IL-4, IL-5, IL-6, IL-7, IL-8 (also known as CXCL8), IL-9, IL-10, IL-12 (p70), IL-13, IL-15, IL-17α, IP10 (CXCL10), MCP1 (CCL2), MIP-1α (CCL3), MIP-1β (CCL4), PDGFB, RANTES (CCL5), TNF-α, VEGF, IL-1α, IL-2Rα, IL-3, IL-12 (p40), IL-16, GRO-α, HGF, IFN-α2, LIF, MCP-3, MIG, β-NGF, SCF, SCGF-β, SDF-1α, CTACK, MIF, TRAIL, IL-18, M-CSF, TNF-β were detected by using the premixed 48-plex Bio-Plex Pro Human Cytokine Assay (Bio-Rad, Hercules, CA, USA; 12007283) according to the manufacturer’s instructions.

### Lymphocyte proliferation assay

For the proliferation analysis, peripheral blood mononuclear cells (PBMCs) from umbilical cord blood of healthy donors were stimulated by 5 µg/mL phytohaemagglutinin (PHA; Sigma-Aldrich, L4144) in a 96-well plate for 3 days in the presence or absence of IMRCs (2 × 10^5^/well) at a ratio of 1:1. At day 3, the proliferation of PBMCs was analyzed by CCK8.

### Real-time quantitative PCR

For reverse transcription (RT), RNA was extracted using an RNAprep Pure Cell/Bacteria Kit (TIANGEN, Beijing, China; DP430). Two micrograms of RNA were reverse transcribed into cDNA using a PrimeScript™ First-Strand cDNA Synthesis Kit (TaKaRa, Shiga, Japan; 6110B). Real-time quantitative PCR (qPCR) was performed and analyzed using a Stratagene Mx3005P system (Agilent Technologies, Santa Clara, CA, USA) with SYBR Green Real-Time PCR Master Mix Plus (Toyobo, Osaka, Japan; QPS-201). *GAPDH* was used for internal normalization. Primers for real-time PCR in this study are as shown in Supplementary information, Table S2.

### Enzyme-linked immunosorbent assay (ELISA)

To determine the secretion of human MMP1, cell culture supernatants were collected after 48 hours of culture. The levels of human MMP1 protein were measured by a Human MMP1 ELISA Kit (Cusabio, Wuhan, China; CSB-E04672h) according to the manufacturer’s instructions. Concentrations of TNF-α (Expandbio, Beijing, China; Exp210030) and TGF-β1 (Expandbio, Exp210089) in the mouse lung homogenates were measured using ELISA kits according to the manufacturer’s instructions.

### Human MMP1 activity

To determine the activity of human MMP1, cell culture supernatants were collected after 48 hours of culture. The activity of human MMP1 were measured by a SensoLyte^®^ Plus 520 MMP-1 Assay Kit (AnaSpec, San Jose, CA, USA; AS-72012) according to the manufacturer’s instructions. A specific anti-MMP-1 monoclonal antibody was used in combination with an MMP1 fluorogenic substrate, 5-FAM/QXL^®^520 FRET peptide. The fluorescence signal is monitored at Ex/Em = 490 nm/520 nm upon MMP-1-induced cleavage of the FRET substrate.

### Immunofluorescence staining

Cells were fixed with 4% paraformaldehyde for 15 min, permeabilized with 0.5% Triton X-100 for 15 min and blocked with 2% BSA for 60 min at room temperature. Then, the cells were stained with 1:200 mouse monoclonal IgG E-cadherin (Abcam, San Francisco, CA, USA; ab1416), 1:200 rabbit monoclonal IgG Collagen I (Cell Signaling Technology, Danvers, MA, USA; 84336), 1:200 rabbit anti-FABP-4 (Thermo Fisher Scientific, PA5-30591), 1:200 rabbit anti-Osteocalcin (Thermo Fisher Scientific, PA5-11849), 1:200 mouse anti Aggrecan (Thermo Fisher Scientific, MA3-16888) overnight at 4 °C in 2% BSA in PBS. The cells were washed 3 times with PBS and then incubated with 1:200 Alexa Fluor 488 donkey anti-mouse IgG (H+L) (Jackson ImmunoResearch, West Grove, PA, USA; 715-545-151), 1:200 Fluorescein (Cy3) donkey anti-rabbit IgG (H + L) (Jackson ImmunoResearch, 711-165-152), 1:200 Fluorescein (Cy3) donkey anti-mouse IgG (H + L) (Jackson ImmunoResearch, 715-165-151) or Fluorescein (Cy2) donkey anti-rabbit IgG (H + L) (Jackson ImmunoResearch, 711-545-152) secondary antibodies in 2% BSA for 60 min at room temperature in the dark. The washing step was repeated before staining the nuclei with Hoechst 33342 (Invitrogen, H3570).

### Western blot

Cells were harvested in RIPA Lysis Buffer (strong) (CWbio, Taizhou, China; CW2333S) containing protease inhibitors (Roche, 4693124001). A total of 40 µg proteins were separated by 4-20% gradient SDS-PAGE gels (GenScript, M42015C) and transferred to PVDF membranes (Millipore, Billerica, MA, USA; IPVH00010). The membranes were blocked at room temperature with 1% BSA for 1 h and incubated overnight at 4 °C with primary antibodies: rabbit Collagen I antibody (Cell Signaling Technology, 84336), mouse E-cadherin antibody (Abcam, ab1416), mouse α-SMA antibody (Sigma-Aldrich, A5228) and mouse β-actin antibody (Sigma-Aldrich, A1978). Then, the membranes were washed with TBST for 3 times and incubated for 1 h with a secondary antibody: anti-mouse IgG antibody (HRD) (Sigma-Aldrich, A9044) and anti-rabbit IgG antibody (HRD) (Sigma-Aldrich, A0545) at room temperature. Images were obtained using the ChemiDoc XRS+ imaging system (Bio-Rad) and quantified using the Quantity One software.

### Soft agar colony formation assay

Soft agarose gels were prepared according to previously published protocols^29^. Cells were harvested and diluted to a cell concentration of 2.5 × 10^4^ cells/mL, before mixing with the agarose mixture in 6-well culture plates. The plates were incubated in a 37 °C incubator for 21 days.

### Animals

Adult female and male cynomolgus monkeys (3-5 years old) were housed in single quarters with a 12/12 h light/dark cycle. Animals were provided with monkey chow and enrichment food twice a day and water *ad libitum*. The monkey experiments were performed at the National Center for Safety Evaluation of Drugs, National Institutes of Food and Drug Control (Beijing, China). The Institutional Animal Care and Use Committee of the National Center for Safety Evaluation of Drugs approved all monkey studies (NO. IACUC-2018-k001).

Specific pathogen-free (SPF) 11- to 13-week-old C57BL/6 male mice were purchased from the Animal Center of Peking University (Beijing, China). All experimental and control mice were weight-matched, and their weights ranged from 25 to 30 g. All mice were housed in the animal care facility of the Institute of Medical Science, China-Japan Friendship Hospital. Mice were maintained under SPF conditions at room temperature (between 20-24 °C), with humidity between 35-55%, in a 12/12 h light/dark cycle, with food and water *ad libitum* and monitored with respect to their general state, fur condition, activity, and weight according to institutional guidelines. The mice were sacrificed at each observational time point by using intraperitoneal pentobarbital overdose. The Animal Studies Committee of the China-Japan Friendship Hospital (Beijing, China) approved all mice studies (NO.190108).

### Acute toxicity and long-term toxicity test

Acute toxicity test: Low dose (2.6 × 10^6^ cells/monkey), middle dose (2.6 × 10^7^ cells/monkey) and high dose (1 × 10^8^ cells/monkey) IMRCs were infused into cynomolgus monkeys (3-5 years old) by single intravenous infusion. After 6 months, their body weight, ophthalmology status, hematology, blood chemistry and urine chemistry were measured.

Long-term toxicity test: Low dose (2.6 × 10^6^ cells/monkey) and high dose (1 × 10^8^ cells/monkey) IMRCs, or saline, were injected into cynomolgus monkeys (3-5 years old) by intravenous infusion once a week for 22 times, after which their body weight, body temperature, food intake and organ weights were measured.

### Copy number variation calling

30X whole-genome sequencing data was produced by HiSeq X-Ten (Annoroad Gene Technology Co., Ltd) and used to analyze single nucleotide mutation changes between IMRCs and hESCs. As previously described^30^, the whole-genome sequencing data was mapped to the hg19 reference genome by BWA (version 0.7.15) using the ‘mem’ mode with default parameters. The genome coverage was calculated by bedtools^31^. The normalized coverage depth for each bin was calculated by dividing the raw coverage depth by the average sequencing depth. Duplicate reads were removed and the uniquely mapped reads were retained for the copy-number variation (CNV) analysis, in which chromosomal sequences were placed into bins of 500 kb in length. The hg19 genome repeat regions annotated by RepeatMasker (http://www.repeatmasker.org) were removed from the genomic sequence before calculating the coverage. The CNV scatterplot was drawn using ggplot2.

### Histological staining and immunostaining staining analysis

Mouse lung histology was performed as follows. Briefly, the lung was dehydrated, paraffin-embedded and cut into 4 µm sections. Lung sections were stained with H&E and Masson’s trichrome stain for assessment of pathological changes. Immunohistochemical staining was performed using antibodies against α-SMA (Servicebio, Wuhan, China; GB13044), Fibronectin (Servicebio, GB13091), Collagen I (Servicebio, GB13091) and GFP (Servicebio, GB13227). Immunofluorescence staining was performed using antibodies against GFP (Servicebio, GB13227), CD31 (Servicebio, GB11063-3) and SPC (Millipore, AB3786).

### Labeling IMRCs with DiR dye

IMRCs were labeled with a lipophilic, near-infrared fluorescent dye 1,1-dioctadecyl-3,3,3,3-tetramethyl indotricarbocyanine iodide (DiR; YEASEN, Shanghai, China; 40757ES25). IMRCs were suspended at a concentration of 1 × 10^6^ cells/mL and incubated with DiR buffer for 20 min at 37 °C according to the manufacturer’s protocol. After incubation, cells were centrifuged at 1000-1500 rpm for 5 min. After washing twice with PBS and resuspension, the IMRCs were preheated at 37 °C before infusion. Both DiR-labeled and unlabeled cells (5 × 10^6^ cells in 200 µL saline) were injected via the tail vein.

### In vivo imaging of IMRCs’ biodistribution

Mice were lightly anesthetized and were monitored using the in vivo imaging systems at day 0, day 1, day 3, day 5, day 7, day 9, day 15, day 21, day 27, day 32, day 37 and day 46 after tail intravenous injection of DiR-labeled IMRCs (5 × 10^6^ cells in 200 µL saline). Serial fluorescence images were also obtained in major organs ex vivo. In order to reduce autofluorescence, the ideal filter conditions for DiR imaging were an excitation/emission spectrum in the near infrared range (750/780 nm)^32,33^.

### Lung function

The flexiVentFX experimental platform was set up according to the manufacturer’s instructions. At the day 21, after 3% pentobarbital sodium (40 mg/kg) intraperitoneal injection of anesthesia, mice were fixed supine within 5-8 minutes, and endotracheal intubation was inserted in the middle of a tracheotomy. After computer processing, indicators of lung function required were obtained, including the inspiratory capacity (IC), respiratory resistance (Rrs), static compliance (Crs), elastic resistance (Ers), Newtonian resistance, tissue damping (G), tissue elasticity (H) and forced vital capacity (FVC)^34^.

### Lung coefficient

The lung tissue was completely removed, weighed by electronic balance, and the lung coefficient was calculated according to the formula: wet lung weight (g)/total body weight (kg).

### Cell therapy in BLM mouse model of pulmonary fibrosis

The BLM mouse model was generated by intratracheal injection of 1.5 mg·kg^-1^ bleomycin sulphate (BLM; Bioway, China; DP721) in normal saline under light anaesthesia. The IMRCs were delivered intravenously at day 0 or day 1 or day 7 or day 14 after injury. Mice were euthanized with 50 mg/mL sodium pentobarbital (0.6 mg/10 g weight) according to published guidelines^35^. Animals were killed at day 21 after BLM injury. After perfusion with normal saline, the left lungs were used for morphometric analysis while the right lungs were removed and used for other analyses.

### Micro-CT

Mouse CT scans were performed according to the manufacturer’s instructions.

### Compassionate use study of two severely ill COVID-19 patients

We conducted a pilot study using GMP-grade IMRCs, under an expanded access program for compassionate use in COVID-19 patients. The pilot study was approved by the Ethics Committee of the Beijing Youan Hospital, Capital Medical University (LL-2020-010-K, Youan EC [2020]005). After two severely ill COVID-19 patients, who tested positive for SARS-CoV-2 by qPCR and were diagnosed with ALI (aged 44-46 years; 1 male and 1 female), gave their consent and met the eligibility criteria, their serum samples were collected and processed in accordance with the ethical legislation and the guidelines of the Ethics Committee of the Beijing Youan Hospital, Capital Medical University.

### Statistics

All data are expressed as mean ± SEM. Survival curves were derived by the Kaplan-Meier method and compared via generalized Wilcoxon test. Statistical analysis was performed using GraphPad Prism 5.0 statistical software (San Diego, CA, USA). The statistical significance of multiple groups was compared to each other using Tukey’s multiple comparison test ANOVA. A p value of < 0.05 was considered statistically significant.

## Results

### Generation of hESC-derived IMRCs

In this study, hESC-derived IMRCs were generated by passaging cells that are migrating from human embryoid bodies (hEBs), using serum-free reagents (Fig. 1a, b). The clinical hESC line (Q-CTS-hESC-2) was prepared as described previously^20^. Clinical hESCs were maintained in Essential 8™ basal medium (E8) on vitronectin-coated plates, before dissociation into small clumps to form hEBs for 5 days. Subsequently, hEBs were transferred onto vitronectin-coated plates and cultured for 14 additional days. The hEBs outgrowth cells were dissociated and passaged continuously in IMRCs Medium. After 5 passages, IMRCs were harvested for characterization. IMRCs possessed fibroblast-like morphology (Fig. 1b) and maintained diploid karyotypes at passage 5 (Fig. 1c). Moreover, copy-number variation analysis by whole genome sequencing data also indicated that no chromosomal aneuploidies, large deletions nor duplication fragments were detected (Fig. 1d). Next, we analyzed the expression profile of MSC-specific genes (Fig. 1e). IMRCs showed a pattern that greatly differed from hESCs, and closely resembled primary umbilical cord-derived MSCs (UCMSCs). All pluripotency genes (*POU5F1, SOX2, NANOG, ZFP42, SALL4*), mesendoderm genes (*MIXL1*), and ectoderm genes (*GAD1*) were extinguished in both IMRCs and UCMSCs (Fig. 1e). IMRCs expressed MSC-specific genes, including *PDGFRA* and *SPARC*, and the MSC-specific surface markers *NT5E* (CD73), *ENG* (CD90), *THY1* (CD105) and *ITGB1* (CD29). Flow cytometry analysis further confirmed this surface marker profile (Fig. 1f; Supplementary information, Fig. S1a, b). By contrast, IMRCs were negative for the hematopoietic surface markers *PTPRC* (CD45) and CD34. IMRCs displayed the ability to undergo tri-lineage differentiation into mesenchymal tissues, such as adipocytes, chondroblasts and osteoblasts (Fig. 1g; Supplementary information, Fig. S1c). The proliferation rate of IMRCs was higher than that of UCMSCs at passage 15, suggesting that IMRCs have a stronger capacity for long-term self-renewal than primary MSCs (Fig. 1h). Interestingly, IMRCs were generally smaller than UCMSCs (Fig. 1i), suggesting that IMRCs can pass through small blood vessels and capillaries more easily, and are thus perhaps less likely to cause pulmonary embolism. To evaluate the clinical potential of the IMRCs, we measured the viability of IMRCs suspended in a published clinical injection buffer at 4 °C. We found that the viability of IMRCs remained higher (93%) than UCMSCs (73%) after 48h (Fig. 1j).

**Fig. 1.**
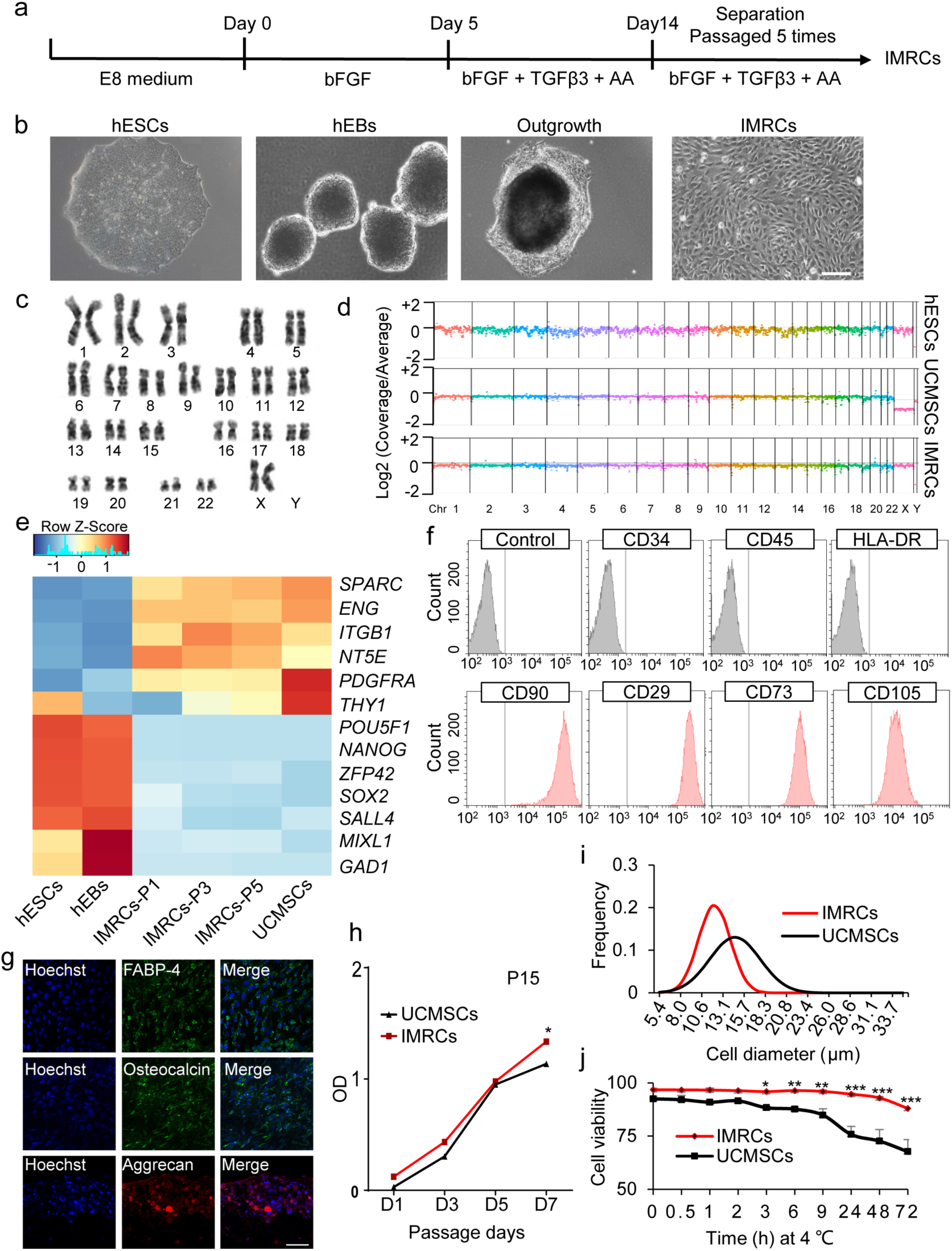
Derivation of immunity- and matrix-regulatory cells (IMRCs) from human embryonic stem cells (hESCs). **a** Different phase of the IMRCs derivation protocol. AA, L-ascorbic acid. **b** Representative morphology of cells at different stages as observed by phase contrast microscopy. hEBs, human embryoid bodies. Scale bars, 100 µm. **c** A representative chromosome spread of normal diploid IMRCs with 22 pairs of autosomes and two X chromosomes. **d** Copy number variation (CNV) analysis by whole genome sequencing for hESCs, primary UCMSCs and IMRCs. UCMSCs, umbilical cord mesenchymal stem cells. **e** Heatmap showing MSC-specific marker and pluripotency marker gene expression changes, from hESCs and hEBs to IMRCs at passages 1-5 (P1-5), and primary UCMSCs. **f** IMRCs’ expression of MSC-specific surface markers was determined by flow cytometry. Isotype control antibodies were used as controls for gating. Like MSCs, the IMRCs are CD34^-^/CD45^-^/HLA-DR^-^/CD90^+^/CD29^+^/CD73^+^/CD105^+^ cells. **g** Representative immunofluorescence staining of IMRCs after they were induced to undergo adipogenic differentiation (FABP-4), osteogenic differentiation (Osteocalcin), and chondrogenic differentiation (Aggrecan). Scale bars, 100 µm. **h** Proliferation curve of IMRCs and UCMSCs at the 15th passage (n = 5). **i** Distribution of cell diameters of IMRCs and UCMSCs. **j** Viability of IMRCs and UCMSCs in clinical injection buffer over time at 4 °C. * *p* < 0.05, ** *p* < 0.01, *** *p* < 0.001; data are represented as the mean ± SEM.

### IMRCs possess unique gene expression characteristics

To clarify the degree of similarity between hESCs, IMRCs and primary UCMSCs at the whole transcriptome level, we performed genome-wide profiling of IMRCs and UCMSCs and compared their gene expression with hESCs^23^. Whole-transcriptome analysis confirmed that IMRCs clustered together with UCMSCs in an unsupervised hierarchical clustering (Fig. 2a). The global differentially expressed gene analysis showed that highly expressed genes in IMRCs and UCMSCs compared with hESCs, were enriched with angiogenesis, inflammatory response and extracellular matrix disassembly related processes (Supplementary information, Fig. S2). Accordingly, MSC-specific genes such as *NT5E, ENG, PDGFRA, SPARC* and *ITGB1* were upregulated, whereas pluripotency genes such as *POU5F1, SOX2, SALL4* and *ZFP42* were extinguished in IMRCs relative to hESCs, and the overall correlation with hESCs was weak (R^2^ = 0.66; Fig. 2b). Next, we analyzed the expression of genes specific to IMRCs, compared to UCMSCs (Fig. 2c). While the overall correlation with UCMSCs was stronger (R^2^ = 0.87), we also found that many genes were differentially expressed in IMRCs compared to primary UCMSCs. The upregulated genes promote immunomodulation (*LIF*), tissue repair (*VEGFA, GREM1*), cell division (*CDC20*) and anti-fibrosis (*MMP1*). By contrast, the downregulated genes predominantly promote inflammation (*IL-1B, CXCL8, CCL2* and *CXCL1;* Fig. 2c). Gene set enrichment analysis (GSEA) of the differentially expressed genes confirmed that IMRCs manifest reduced inflammation and stronger proliferative capacity as their top gene signatures, compared to primary UCMSCs (Fig. 2d, e; Supplementary information, Fig. S3).

**Fig. 2.**
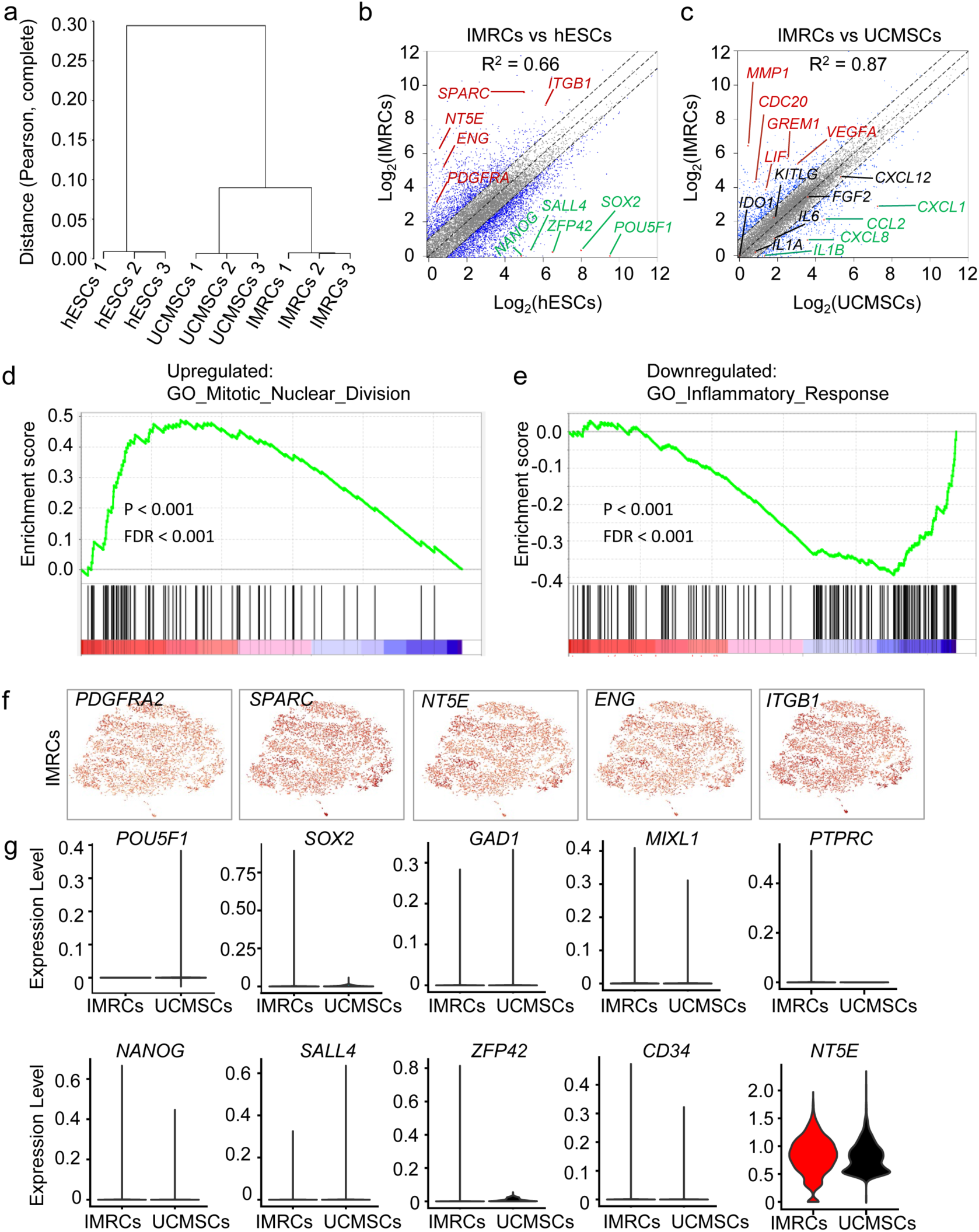
IMRCs possess unique gene expression characteristics. **a** Unsupervised hierarchical clustering analysis based on the Pearson correlation distance between the whole mRNA profile of each cell type. **b** Scatter plot displaying the differentially expressed genes (DEGs) between IMRCs and hESCs. Up-regulated genes are highlighted in red. Down-regulated genes are highlighted in green. Gray dots represent non-DEGs (< 2-fold change). **c** Scatter plot displaying the DEGs between IMRCs and primary UCMSCs. Up-regulated genes are highlighted in red. Down-regulated genes are highlighted in green. Gray dots represent non-DEGs (< 2-fold change). **d** Gene set enrichment analysis (GSEA) of the top up-regulated gene signature in IMRCs, compared with primary UCMSCs. **e** GSEA of the top down-regulated gene signature in IMRCs, compared with UCMSCs. **f** Heatmaps of specific gene expression amongst single IMRCs groups. **g** Quantification of non-mesenchymal marker gene expression amongst single IMRCs, UCMSCs and hESCs, as measured by single cell RNAseq.

To elucidate the heterogeneity of gene expression amongst IMRCs, single cell RNA sequencing (scRNAseq) was performed for both IMRCs and primary UCMSCs. A total of 16,000 single-cell transcriptomes were obtained from two samples. Approximately ∼ 100,000 reads were obtained per cell, which generated a median of 31,000 unique molecular identifiers per cell, ∼ 4,800 expressed genes per cell and more than 22,000 total genes detected in the population. Our scRNAseq data also showed that IMRCs were relatively homogenous in expression for the MSC-specific markers *PDGFRA, SPARC, NT5E, ITGB1* and *ENG* (Fig. 2f; Supplementary information, Fig. S4). IMRCs were also relatively homogenous in their suppression of pluripotency and non-mesenchymal markers, relative to hESCs and UCMSCs, suggesting they are likely to be similar in biological activity to primary MSCs after transplantation (Fig. 2g; Supplementary information, Fig. S4). These results provided insight into the clinical applicability of IMRCs, especially with regards to their complete loss of pluripotency and their gain in hyper-immunomodulatory potential.

### IMRCs treated with IFN-γ show hyper-immunomodulatory potency

To test the immunomodulatory capacity of IMRCs, we exposed them to the pro-inflammatory cytokine IFN-γ. We found that after stimulation with IFN-γ, both primary UCMSCs and IMRCs displayed similar characteristic morphological changes (Fig. 3a). IFN-γ stimulation also potently upregulated *IDO1* (indoleamine 2, 3-dioxygenase 1) expression in both primary UCMSCs and IMRCs, but much more significantly so in IMRCs (Fig. 3b). This is important because *IDO1* has been shown to mediate immunosuppression in T cell-inflamed microenvironments through its catabolism of tryptophan and thus suppression of the tryptophan-kynurenine-aryl hydrocarbon receptor (Trp-Kyn-AhR) pathway in T cells. To further characterize IMRCs at a molecular level, we conducted genome-wide RNA profiling to compare IMRCs with UCMSCs, after IFN-γ stimulation (Fig. 3c). Hierarchical clustering revealed that IMRCs clustered separately from UCMSCs, although they appeared more similar after IFN-γ stimulation (Fig. 3d). To analyze IMRCs after IFN-γ stimulation in greater detail, we separated their differentially expressed genes into different categories and tested their overlap using Venn diagram analysis (Fig. 3e). The global differential expressed gene analysis showed that highly expressed genes in IMRCs and UCMSCs after IFN-γ stimulation, compared with IMRCs and UCMSCs before IFN-γ stimulation, were enriched with immune response and interferon-gamma-mediated signaling pathway (Supplementary information, Fig. S5). Accordingly, we found that some pro-inflammatory genes showed lower expression while some pro-regenerative genes showed higher expression levels in IMRCs, compared to primary UCMSCs (Fig. 3f).

**Fig. 3.**
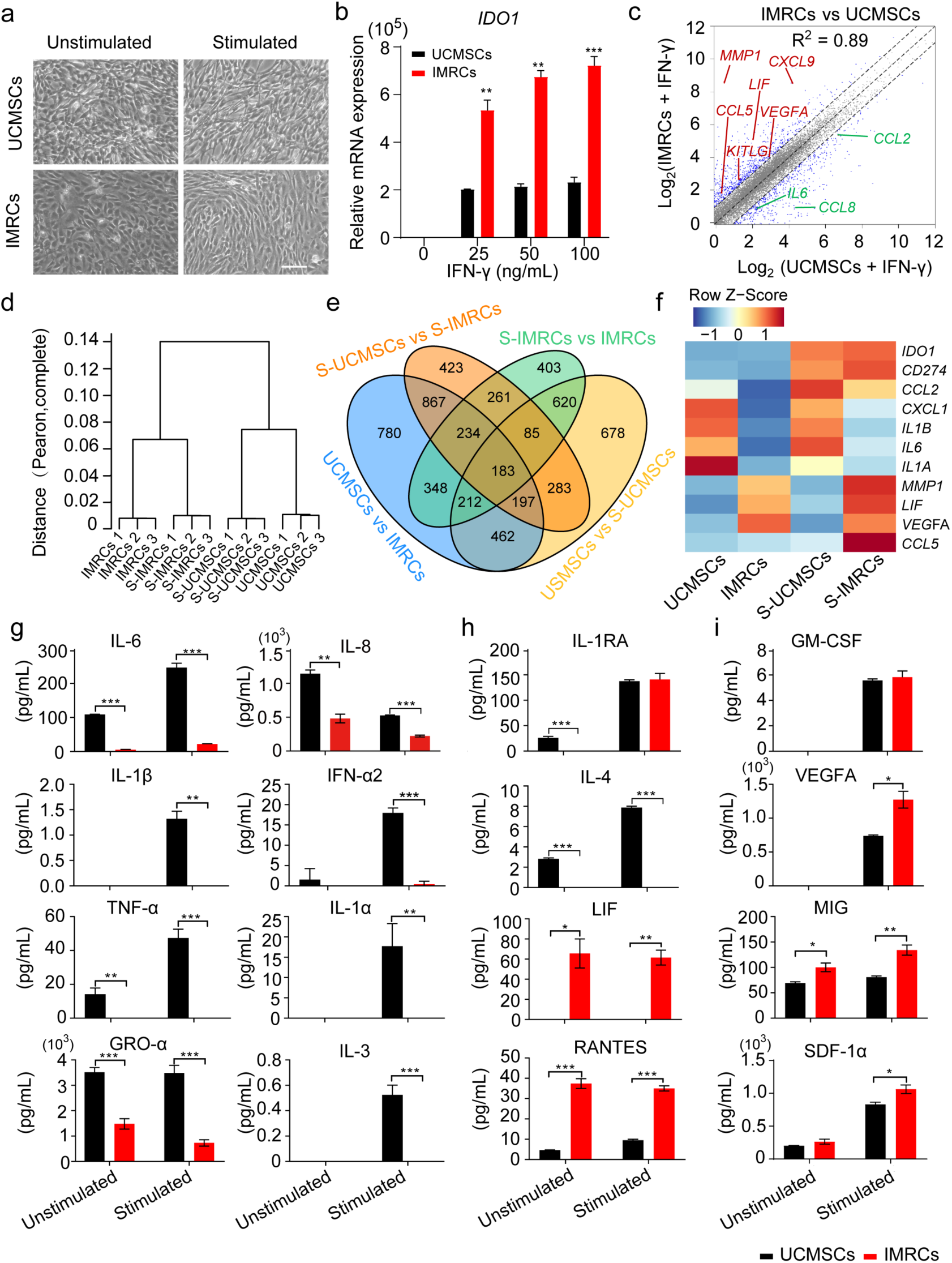
IMRCs activated by IFN-γ show hyper-immunomodulatory potency. **a** Morphology of UCMSCs and IMRCs before and after IFN-γ (100 ng/mL) stimulation. **b** Quantitative PCR for *IDO1* mRNA in UCMSCs and IMRCs before and after IFN-γ (0, 25, 50, 100 ng/mL) treatment. *GAPDH* was used for internal normalization. **c** Scatter plot displaying the DEGs between UCMSCs and IMRCs after treatment with 100 ng/mL IFN-γ. Up-regulated genes are highlighted in red. Down-regulated genes are highlighted in green. Gray dots represent non-DEGs (< 2-fold change). **d** Unsupervised hierarchical clustering analysis of UCMSCs and IMRCs, before and after IFN-γ stimulation (S-UCMSCs and S-IMRCs). **e** Venn diagram shows the overlap among UCMSC- and IMRCs-associated genes, before and after IFN-γ stimulation (S). **f** Heatmap illustrating the expression of cytokines in UCMSCs and IMRCs, before and after IFN-γ stimulation (S). **g-i** ELISA analysis of biologically relevant chemokines and cytokines in the secretomes of unstimulated or stimulated IMRCs and UCMSCs. Pro-inflammatory (G), immunomodulatory (H), and pro-regenerative (I) cytokines. * *p* < 0.05, ** *p* < 0.01, *** *p* < 0.001; data are represented as the mean ± SEM. Scale bar, 100 µm.

To confirm these findings, we performed a focused ELISA analysis of 48 biologically relevant chemokines and cytokines in the secretomes of both stimulated IMRCs and UCMSCs. We found that at least nine pro-inflammatory cytokines were lower in IMRCs than UCMSCs, including interleukin-6 (IL-6), IL-1β, tumor necrosis factor alpha (TNF-*α*), GRO-*α*, IFN-*α*2, IL-1*α*, IL-3, and IL-8 (Fig. 3g; Supplementary information, Fig. S6a). Furthermore, while the anti-inflammatory IL-4 was lower, the immunomodulatory cytokines IL-1RA, LIF, and RANTES were higher in IMRCs than UCMSCs (Fig. 3h). Amongst the pro-regenerative cytokines, we found that GM-CSF was similar while VEGFA, MIG and SDF-1α were higher in IMRCs than UCMSCs (Fig. 3i). To determine the immunomodulatory properties of IMRCs, we examined the effects of IMRCs on PBMCs’ proliferation and found that IMRCs significantly inhibited PHA-stimulated PBMCs’ proliferation when cocultured together at a ratio of 1:1 (Supplementary information, Fig. S6b). These results indicate that IMRCs manifest much stronger immunomodulatory and pro-regenerative functions than UCMSCs.

### IMRCs suppress the pro-fibrotic effects of TGF-β1

Our scRNAseq results also indicated that more than 99% of IMRCs expressed *MMP1* compared with primary UCMSCs (< 1%) (Fig. 4a; Supplementary information, Fig. S4). Similarly, gene expression analysis by RT-qPCR showed that IMRCs express much higher levels of *MMP1* than hESCs, UCMSCs or human foreskin fibroblasts (HFF; Fig. 4b). MMP1 could be detected as a highly secreted protein in the conditioned media of IMRCs by ELISA (Fig. 4c). In addition, the secreted MMP1 was an enzymatically active and correctly processed isoform (Fig. 4d, e). To investigate the role of the IMRCs on fibrosis, the human epithelial cell line A549 was used to establish an in vitro fibrosis model. The morphology of A549 cells became myofibroblast-like after 48 h treatment with 10 ng/mL TGF-β1 (Fig. 4f). Consistent with the morphological changes, the expression of myofibroblast markers such as *ACTA2, Collagen I, Collagen II, Fibronectin* and *TGFB1* increased, while the expression of the epithelial identity marker *CDH1* was extinguished, suggesting that the A549 cells had undergone TGF-β1-induced myofibroblast transdifferentiation (Fig. 4g). Immunofluorescence and western blot analyses confirmed that TGF-β1 induced the upregulation of collagen I and the downregulation of E-cadherin (Fig. 4h, i). These results indicated that the A549 alveolar epithelial cells could simulate lung fibrosis in vitro. Thereafter, A549 cells were cultured with or without the conditioned media of IMRCs, along with TGF-β1 treatment. Our results showed that the IMRC conditioned media significantly ameliorated the induction of *Collagen I, ACTA2* and *TGFB1* expression (Fig. 4j; Supplementary information, Fig. S7a). Immunofluorescence and western blot analyses further confirmed that IMRC conditioned media could extinguish Collagen I protein expression and inhibit α-SMA protein expression during the A549-to-myofibroblast transdifferentiation (Fig. 4k-m; Supplementary information, Fig. S7b, c). These results suggested that IMRCs might be able to inhibit lung fibrosis.

**Fig. 4.**
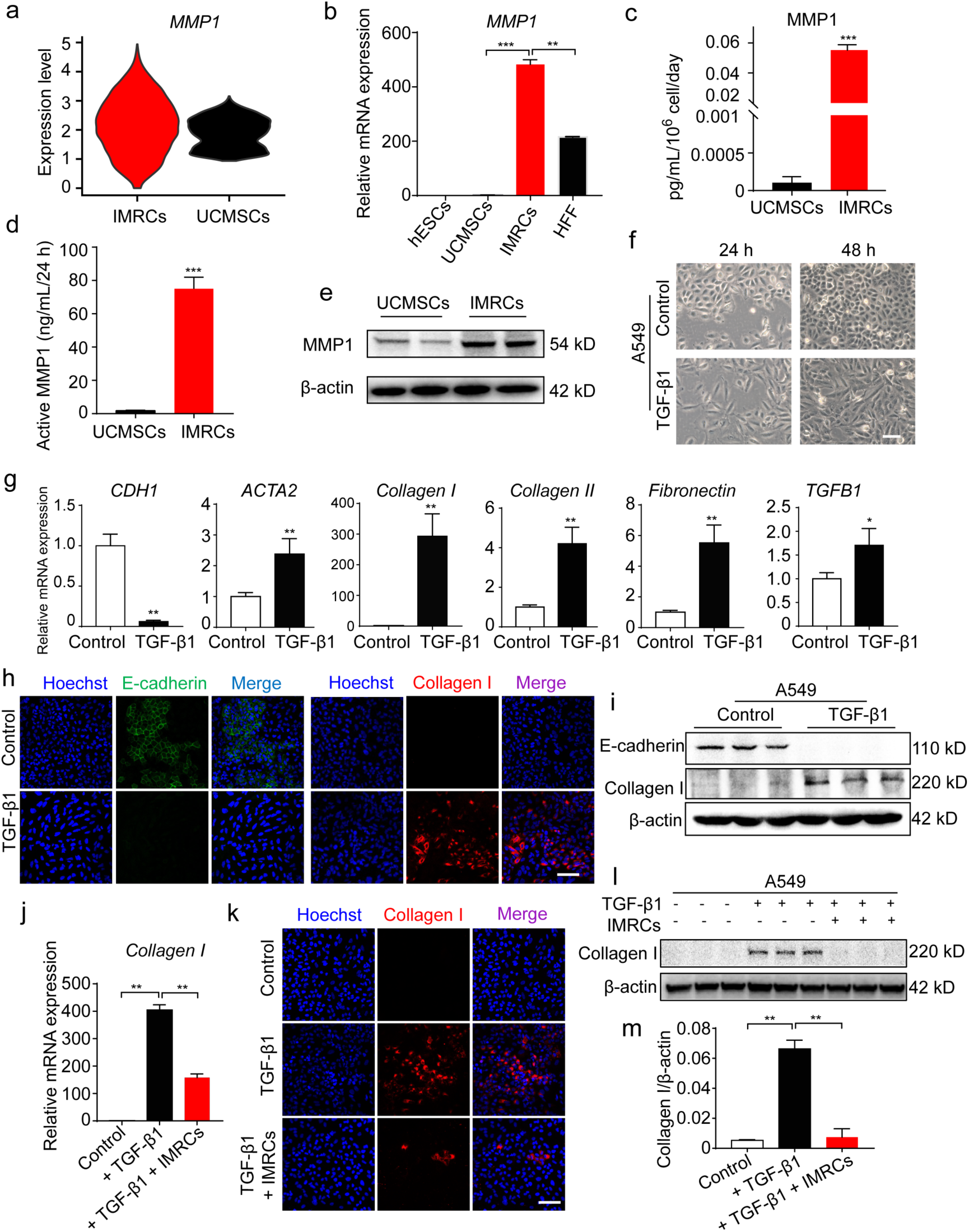
IMRCs reduce the pro-fibrotic effects of TGF-β1. **a** Quantification of *MMP1* gene expression amongst single IMRCs, UCMSCs and hESCs, as measured by single cell RNAseq. **b** Quantitative PCR for *MMP1* mRNA in hESCs, UCMSCs, IMRCs and human foreskin fibroblasts (HFF). **c** ELISA for MMP1 protein in the conditioned media of IMRCs and UCMSCs. **d** MMP1 activity in the conditioned media of IMRCs and UCMSCs. **e** Western blot for MMP1 protein in IMRCs and UCMSCs. β-actin was used as a loading control. **f** Representative morphology of A549 cells, with or without 10 ng/mL TGF-β1 treatment for 48 h. **g** Quantitative PCR for the relative expression of *CDH1, ACTA2, Collagen I, Collagen II, Fibronectin* and *TGFB1* mRNA in A549 cells, with or without TGF-β1 treatment for 48 h. **h** Immunofluorescence staining for E-cadherin and Collagen I expression in A549 cells, with or without 10 ng/mL TGF-β1 treatment for 48 h. **i** Western blot for E-cadherin and Collagen I protein expression in A549 cells, with or without 10 ng/mL TGF-β1 treatment for 48h. **j** Quantitative PCR for *Collagen I* mRNA in A549 cells, with or without 10 ng/mL TGF-β1 and IMRC conditioned media treatment for 48 h. **k** Immunofluorescence staining for E-cadherin and Collagen I expression in A549 cells, with or without 10 ng/mL TGF-β1 and IMRC conditioned media treatment for 48 h. **l** Western blot for E-cadherin and Collagen I protein expression in A549 cells, with or without 10 ng/mL TGF-β1 and IMRC conditioned media treatment for 48 h. **m** Quantification of the relative Collagen I protein expression levels in **(l)**. * *p* < 0.05, ** *p* < 0.01, *** *p* < 0.001; data are represented as the mean ± SEM. Scale bar, 100 µm.

### Safety evaluation of IMRC transfusion into mice and monkeys

Tumorigenicity has long been an obstacle to the clinical application of cells derived from hESCs, due to the contamination of residual hESCs that could form teratomas. Our single cell RNA sequencing showed that no residual OCT4^+^/SOX2^+^/NANOG^+^/TERT^+^/DPPA5^+^ hESCs remained amongst the IMRCs, and that all pluripotency gene expression had been extinguished (Fig. 5a). To ensure their short-term and long-term safety, a series of biosafety-related experiments were performed according to the “Guidelines for Human Somatic Cell Therapies and Quality Control of Cell-based Products” of China Food and Drug Administration (CFDA) (http://samr.cfda.gov.cn/WS01/CL0237/15709.html), whereupon the IMRCs were verified as suitable for use in human therapy (Supplementary information, Table S1). These biosafety-related experiments included testing for bacteria, fungi, mycoplasma, virus (by in vivo and in vitro methods), pluripotent cell residuals, tumorigenicity and biopreparation safety (endotoxin and bovine serum albumin residuals). According to these tests, the safety of the IMRCs has been verified as required by the National Institutes for Food and Drug Control (NIFDC) in China. To track the biodistribution and long-term engraftment of IMRCs in vivo, whole-animal imaging of wildtype mice was performed at designated time points (day 0, 1, 3, 5, 7, 9, 15, 21, 27, 32, 37 and 46) after injection with DiR-labeled IMRCs (Supplementary information, Fig. S8a). Our data showed that DiR fluorescence intensity declined by half on the 5th day, and dropped steadily over at least 15 days, disappearing around day 46 (Fig. 5b, c; Supplementary information, Fig. S8c). No DiR fluorescence signals were observed in the control mouse. These results indicate that IMRCs never show long-term engraftment in vivo. However, IMRCs were predominantly distributed to the lung, with small amounts in the liver and spleen (Supplementary information, Fig. S8b). To assess the recruitment of IMRCs to the lung in greater detail, GFP-labeled IMRCs were used to track the distribution of IMRCs. A number of GFP-labeled IMRCs were observed in the interstitial spaces in the lung, but not in the lung capillaries (Fig. 5d). Immunofluorescence analysis of the expression of the alveolar type II epithelial cell marker SPC and the endothelial cell marker CD31 showed that neither marker was co-expressed with GFP-labeled IMRCs in the lungs of mice by day 21 (Fig. 5d). These data suggested that IMRCs are unlikely to engraft nor transdifferentiate into endothelial or epithelial cells, after homing to the interstitium of lung tissues in vivo.

**Fig. 5.**
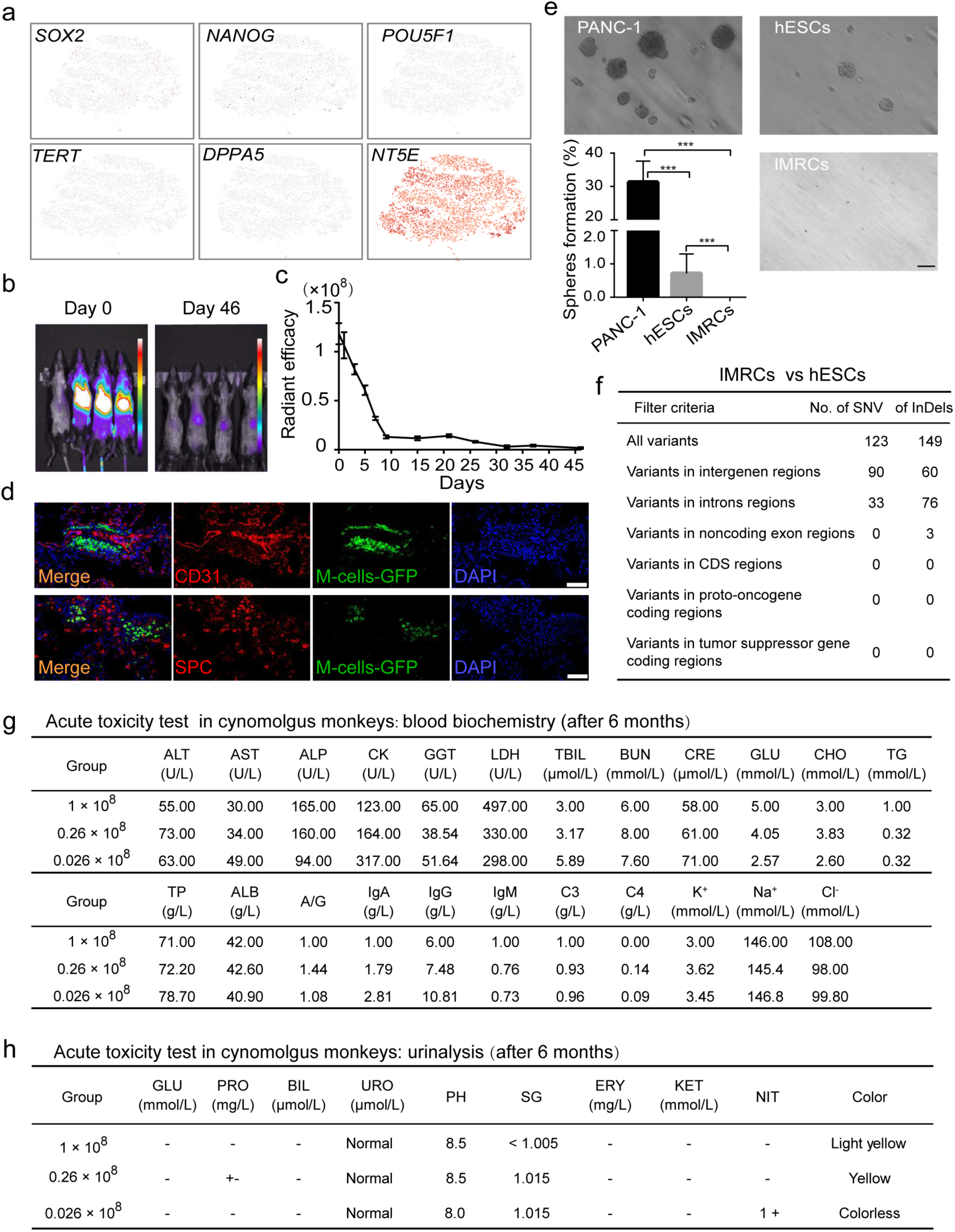
Evaluation of the safety of IMRC transfusion. **a** t-SNE projection of single cells and their pluripotency gene expression amongst IMRCs. **b** In vivo imaging of the far red fluorescence in mice after injection of DiR-labeled IMRCs. **c** Quantification of the DiR fluorescent radiance in the whole body over time. **d** Immunofluorescence staining of lung sections for the endothelial marker CD31 and the alveolar epithelial marker SPC, 21 days after injecting GFP-labeled IMRCs. Scale bar, 100 µm. **e** Soft agar colony formation assay for the tumorigenic potential of IMRCs, relative to PANC-1 cancer cells and hESCs. Scale bar, 100 µm. **f** Single nucleotide variants (SNVs) and insertions/deletions (InDels) in the non-repeat regions of the IMRCs genome, relative to hESCs. **g** Blood biochemistry results of cynomolgus monkeys (*Macaca fascicularis*) injected with a low (2.6 × 10^6^), medium (2.6 × 10^7^) or high (1 × 10^8^) dose of IMRCs after six months. AST, aspartate aminotransferase; ALT, alanine aminotransferase; ALP, alkaline phosphatase; CK, creatine phosphokinase; GGT, glutamyltransferase; LDH, lactate dehydrogenase; TBIL, total bilirubin; BUN, urea nitrogen; CRE, creatinine; GLU, glucose; CHO, total cholesterol; TG, triglyceride; TP, total protein; ALB, albumin; A/G, albumin/globulin. **h** Urinalysis results of cynomolgus monkeys (*Macaca fascicularis*) injected with a low (2.6 × 10^6^), medium (2.6 × 10^7^) or high (1 × 10^8^) dose of IMRCs after six months. PRO, urine protein; BIL, urine bilirubin; URO, urinary gallbladder; pH, acid degree value; SG, specific gravity; ERY, erythrocyte; KET, ketone; NIT, nitrite. * *p* < 0.05, ** *p* < 0.01, *** *p* < 0.001; data are represented as the mean ± SEM.

To give some indication of the tumorigenic potential of IMRCs in vitro, a soft agar assay was performed (Fig. 5e). As a positive control, the PANC-1 pancreas tumor cell line showed a colony formation rate of about 30%. A small number of clones were also formed by hESCs, and the colony formation rate was about 0.5%. However, no colonies were formed by IMRCs. These results indicated that IMRCs had no tumorigenic potential as determined by this assay. Somatic mutation analysis by whole genome sequencing also showed zero mutations in all coding and noncoding exon regions of the IMRCs genome (Fig. 5f). Tumor formation assays also confirmed that IMRCs could not form any tumors in immunodeficient mice after injection in vivo (Supplementary information, Table S1).

We also transfused different doses of IMRCs into cynomolgus monkeys (*Macaca fascicularis*), to evaluate the short- and long-term toxicity of IMRCs in primates. After 6 months, acute toxicity data showed that all blood biochemistry and urinalysis markers remained in the normal range (Fig. 5g, h; Supplementary information, Fig. S9a), indicating normal liver, kidney, heart, muscular, pancreatic and overall metabolic functions. Moreover, no abnormalities were observed in the long-term toxicity tests either, suggesting minimal xenogeneic rejection responses (Supplementary information, Fig. S9b; n = 18). These results indicated that IMRCs have a robust safety profile in vitro and in vivo, and could potentially provide therapeutic treatments with good safety levels for clinical potential.

### IMRC transfusion treats lung injury and fibrosis in a dose-dependent manner

To evaluate the therapeutic effects of IMRCs on lung injury and fibrosis, IMRCs were administered intravenously into a bleomycin-induced model of lung injury (Fig. 6a). IMRCs ameliorated the total body weight reduction in mice exposed to bleomycin (BLM)-induced lung injury, in a dose-dependent manner (Fig. 6b). Kaplan-Meier survival curves indicated that IMRC treatment prolonged the overall survival rates (BLM group 12.5% vs. 1 × 10^6^ IMRCs 25%, 3 × 10^6^ IMRCs 50%, 5 × 10^6^ IMRCs 62.5% (*P* < 0.01)) and the median survival time (BLM group 11.5 d vs. 1 × 10^6^ IMRCs 15.5 d, 3 × 10^6^ IMRCs 18.0 d, 5 × 10^6^ IMRCs 21.0 d) in mice exposed to bleomycin-induced lung injury. There was a statistically significant difference between the BLM group and the 5 × 10^6^ IMRCs group, but not the other groups (Fig. 6c). Histological staining of the lung at day 21 after bleomycin injection (inflammation phase) showed diffuse pneumonic lesions with loss of the normal alveolar architecture, septal thickening, enlarged alveoli, and increased infiltration of inflammatory cells in the interstitial and peribronchiolar areas, in the BLM lung compared with normal lung. IMRC treatment reduced alveolar thickening in the lung in a dose-dependent manner (Fig. 6d-f). Moreover, IMRC treatment reduced the number of macrophages in the lung (Fig. 6f). These results indicated that IMRC treatment can reduce inflammation in the lung after acute injury. Moreover, IMRC treatment also improved the Ashcroft score for pulmonary fibrosis in a dose-dependent manner (Fig. 6g). In particularly, IMRC treatment decreased collagen deposition in the BLM lung in a dose-dependent manner (Supplementary information, Fig. S10b, f). The expression levels of COL I, FN and α-SMA were also significant lower after IMRC treatment (Supplementary information, Fig. S10). ELISA showed that IMRC treatment reduced both TNF-α and TGF-β1 levels in the BLM lung in a dose-dependent manner (Fig. 6h, i). These results suggested that IMRCs can significantly reduce inflammation and fibrosis after lung injury, in a dose-dependent manner.

**Fig. 6.**
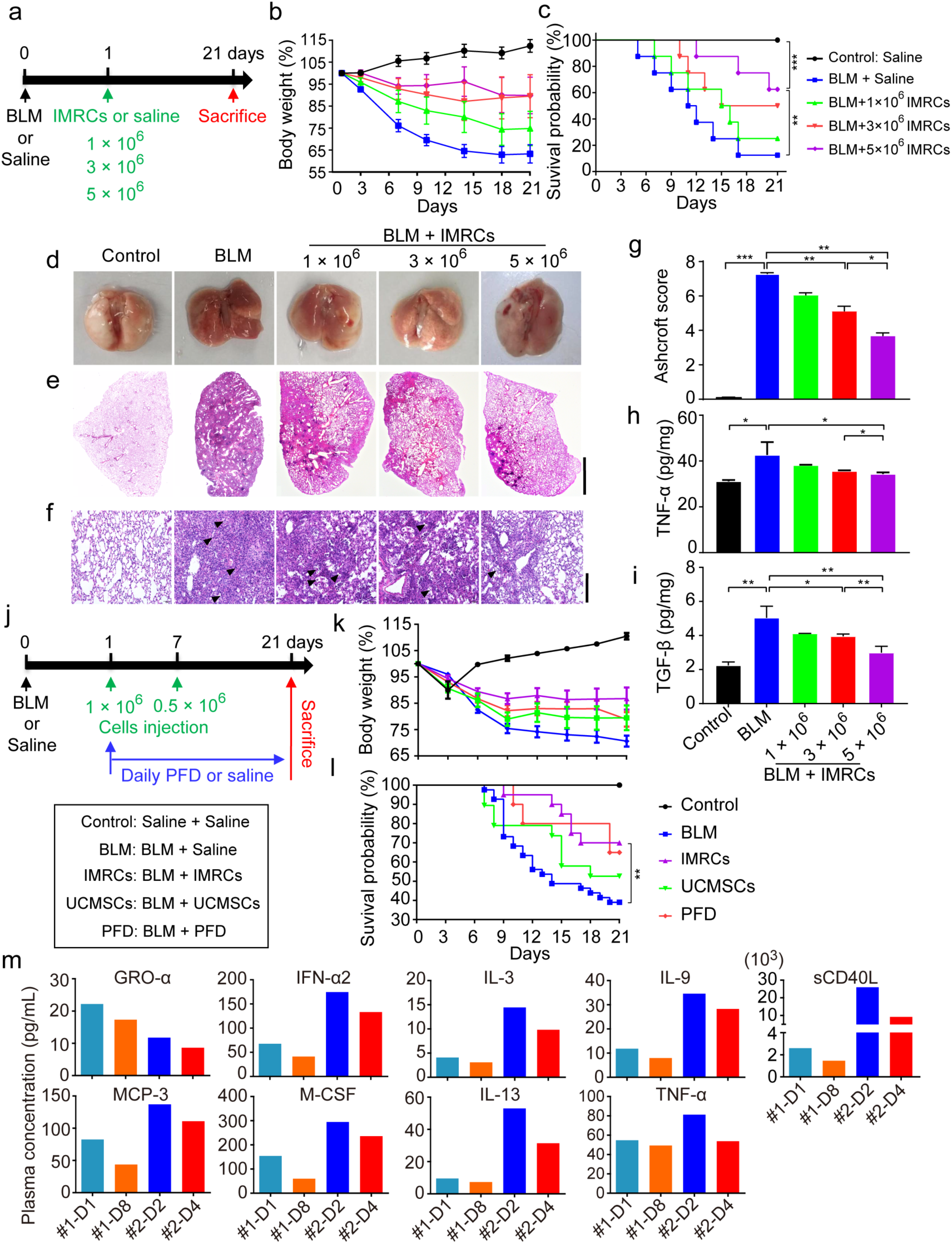
IMRC transfusion treats lung injury and fibrosis dose-dependently. **a** Diagram of the animal experimental protocol for dose escalation. Mice received intratracheal bleomycin (BLM; 2.5 mg/kg body weight) or the same amount of saline at day 0. At day 1, some BLM-injured mice received an intravenous (I.V.) injection of 1 × 10^6^, 3 × 10^6^ or 5 × 10^6^ IMRCs via the caudal vein. A group of BLM-injured mice and normal control mice received the same volume of saline. Mice were randomly grouped (n = 8 per group). **b** Relative body weight (%) changes of the mice receiving different interventions. **c** Kaplan-Meier survival curves of the mice receiving different interventions. **d** Representative images of whole lung from all groups at day 21 post-injury. **e, f** Representative histology of lung sections stained with H&E (Scale bar, 2 mm and 200 µm) at 21 days post-injury. Arrowheads, inflammatory infiltration. **g** Quantitative evaluation of fibrotic changes with the Ashcroft score in lungs of mice receiving different interventions. The Ashcroft scores based on the lung H&E sections. The severity of fibrotic changes in each section was assessed as the mean score of severity in the observed microscopic fields. Six fields per section were analyzed. **h, i** ELISA for the protein levels of TNF-α and TGF-β1 in the lungs of mice receiving different interventions. **j** Diagram of the animal experimental protocol for multiple arm testing. Double I.V. injections of 1 × 10^6^ and 0.5 × 10^6^ IMRCs (at days 1 and 7 respectively) were compared side-by-side with double I.V. injections of 1 × 10^6^ and 0.5 × 10^6^ UCMSCs (at days 1 and 7 respectively), vs. daily pirfenidone (PFD) or saline treatments of mice injured with intratracheal bleomycin (BLM; 2.5 mg/kg body weight) at day 0. **k** Relative body weight (%) changes of mice receiving different interventions. **l** Kaplan-Meier survival curves of mice receiving different interventions. **m** ELISA analysis of pro-inflammatory cytokines in the plasma of two ALI patients after intravenous IMRC transfusion. * *p* < 0.05, ** *p* < 0.01, *** *p* < 0.001; data are represented as the mean ± SEM.

### IMRC treatment of lung injury and fibrosis is superior to UCMSCs and pirfenidone injections

Pirfenidone (PFD) is an FDA-approved drug for the treatment of idiopathic pulmonary fibrosis. It works by reducing lung fibrosis through downregulating the production of growth factors and procollagens I and II. To evaluate the efficacy of IMRC treatment, we compared it side-by-side with UCMSCs and PFD treatments of BLM-injured mice (Fig. 6j). Our results showed that IMRC treatment led to an improved amelioration of the weight reduction after BLM-induced lung injury, compared to UCMSCs or PFD treatments (Fig. 6k). Kaplan-Meier survival curves indicated that the IMRC treatments also resulted in the highest overall survival rates compared to UCMSCs or PFD treatments (BLM group 39.02%, vs. IMRCs 70% (*P* < 0.01), UCMSCs 52.63%, PFD 65% (*P* < 0.05); Fig. 6l). Moreover, IMRC treatment showed the best improvement in lung morphology (Supplementary information, Fig. S11a) and the best reduction in BLM-induced edema, as measured by the lung coefficient (lung wet weight/total body weight), compared to UCMSCs or PFD treatments (Supplementary information, Fig. S11b).

Functionally, mice treated with IMRCs showed the best improvement in indices for lung capacity, including pressure-volume (PV), inspiratory capacity (IC), static compliance (Crs) and forced vital capacity (FVC) (Supplementary information, Fig. S12a-d). IMRC treatment also showed the best reduction in functional indices for lung fibrosis, including respiratory resistance (Rrs), elastic resistance (Ers), tissue damping (G) and tissue elasticity (H) were observed (Supplementary information, Fig. S12e-h). Computed tomography (CT) scans also showed that IMRC treatment led to the best improvement in lung morphology, compared to UCMSCs or PFD treatments (Supplementary information, Fig. S12i, j). In fact, measurements of the lung volume indicated that mice treated with IMRCs had the best improvement in lung volume compared to UCMSCs or PFD treatments. These results indicated that the IMRC therapy is superior to primary UCMSCs and PDF in treating lung injury and fibrosis.

Encouraged by these results, and in response to the urgency of the COVID-19 crisis in China, we pilot-tested IMRC transfusion as compassionate treatments in two severely ill COVID-19 patients who were diagnosed with ALI, as part of an expanded access program. After patient consent, IMRCs were administered intravenously in both severely ill patients. Within 14 days after IMRC transfusion, both the severely ill COVID-19 patients showed significant recovery from pneumonia, tested negative for SARS-CoV-2, and were recommended for discharge. Cytokine analysis of both patients’ plasma showed very high levels of inflammatory cytokines initially in days 1-2, as expected of the ALI-induced cytokine release syndrome that was manifesting at the point of treatment (Fig. 6m). However, by days 4-8, many pro-inflammatory cytokines were suppressed after IMRC infusion, including GRO-*α*, IFN-*α*2, IL-3, IL-9, IL-13, MCP-3, M-CSF, sCD40L and TNF-*α* (Fig. 6m). These results suggested that IMRCs might possess a hyper-immunomodulatory function with potential therapeutic benefits to ALI patients without other treatment options.

## Discussion

In this study, IMRCs were derived from self-renewing hESC cultures with serum-free reagents. Our results showed that IMRCs, while similar to primary UCMSCs, were superior in their long-term proliferative capacity, hyper-immunomodulatory and anti-fibrotic functions. In addition, the cell diameters of IMRCs were generally smaller than UCMSCs, suggesting that they pose lower risks for pulmonary embolism after injection. Detailed analysis of mice and monkeys after IMRC injections indicates that IMRCs do not engraft, nor transdifferentiate nor initiate tumorigenesis, showing excellent potential for short- and long-term safety profiles by a range of in vitro and in vivo assays. Most importantly, our experiments with the bleomycin mouse model of lung injury showed that treatments with IMRCs are superior to primary UCMSCs and the FDA-approved pirfenidone in therapeutic efficacy.

In recent years, MSC-like cells have been generated from hESCs using various protocols^18,19,36,37^. However, to the best of our knowledge and in spite of significant efforts by numerous groups, previously reported studies lacked biosafety-related experiments. This has hindered their translation to clinical application. To ensure safety, a series of biosafety-related experiments were performed according to the “Guidelines for Human Somatic Cell Therapies and Quality Control of Cell-based Products” and the IMRCs were verified as suitable for use in human therapy. Our data showed that there was no teratoma formation observed after IMRC injection into the testes of NOD-SCID mice. In addition, single cell RNA sequencing data showed no residual hESCs were detected in IMRCs and soft agar assays also showed that no colonies were formed by IMRCs. In addition, the somatic mutation analysis by whole genome sequencing also showed no mutations in proto-oncogenes and tumor suppressor genes, nor any coding and non-coding exon regions of IMRCs. These results indicated that IMRCs have no demonstrable potential to form tumors. Moreover, IMRCs could not undergo long-term engraftment and disappeared around day 46. IMRCs did not transdifferentiate into endothelial cells nor alveolar epithelial cells in the lung. To evaluate the risks of clinical application, short- and long-term toxicity tests were performed in cynomolgus monkeys. After 6 months, both acute toxicity tests and long-term toxicity test data showed that no abnormalities were observed. These results proved that IMRCs could have an excellent safety profile and clinical potential. Moreover, the above data suggested that the therapeutic efficacy of IMRCs may not be due to long-term engraftment and direct repair of tissue, but due to their hyper-immunomodulatory and anti-fibrotic functions at inflammatory sites of tissue interstitium.

Like MSCs, the IMRCs are immunoprivileged, and can escape allogeneic immune responses through their lack of expression of HLA class II and their weak expression of HLA class I, thus making them valuable choices for allogeneic cell therapy. Moreover, it has been reported that MSCs could release immunomodulatory factors, alter the expression of surface molecules, and produce growth factors during an immune response to the inflammatory cytokines produced by T cells and antigen-presenting cells^38^. These immunomodulatory factors are crucial for regulating the immune system and promoting tissue repair. Accordingly, expression of the T cell-suppressive *IDO1* greatly increased in IMRCs after stimulation with the pro-inflammatory cytokine IFN-γ, even more than primary MSCs. Genome-wide RNA sequencing revealed that IMRCs and UCMSCs had different expression profiles after exposure to IFN-γ. Analysis of the secretomes of IMRCs and UCMSCs confirmed at least 9 pro-inflammatory cytokines that were lower, and several anti-inflammatory cytokines and pro-regenerative factors that were higher in IMRCs than UCMSCs. All these results suggested that IMRCs possess a hyper-immunomodulatory capacity compared to primary UCMSCs, after stimulation with IFN-γ. Thus, these cells can be administered to immunocompetent animals and human patients without the need for further immunosuppression.

Moreover, higher levels of MMP1 were secreted by IMRCs than primary UCMSCs. MMP1 plays a very important role in the process of fibrosis, by degrading existing collagens and promoting the early stages of tissue remodeling that are critical for the progression of fibrogenesis^39,40^. In general, an imbalance between MMPs and TIMPs is the direct cause of fibrosis and tissue scarring. It has been reported that transplantation of human *MMP1*-overexpressing bone marrow-derived mesenchymal stem cells can attenuate CCL4-induced liver fibrosis in rats^41^. Therefore, it is conceivable that IMRCs that highly express MMP1 may also reverse lung fibrosis. Accordingly, our data showed that the secretomes from IMRCs could reduce collagen I levels during fibrogenesis induced by TGF-β1. However, it is still unclear whether MMP1 played a direct role in the decrease of collagen I. Future work will focus on the role of MMP1 by using *MMP1*-knockout IMRCs in vitro and in vivo.

The mouse BLM model is similar to human ALI/ARDS during the acute inflammatory phase. Although fibroblastic foci, alveolar epithelial type 2 cells hyperplasia and honeycombing lesions are reduced compared with those in humans, indicating that the modeling of human ARDS is not complete in the mouse BLM model, injury to alveolar epithelial cells has been shown to be a common contributing factor to the pathogenesis of both human ARDS and BLM-induced mouse pulmonary injury^41,42^. To test the therapeutic efficacy of IMRCs in the BLM mouse model, the delivery route and schedule of administration (e.g. single dose versus repeated doses) were carefully considered. Intravenous infusion is one of the principal delivery routes, considered to be minimally invasive, simple to use, and is the most common mode for primary MSCs delivery in diverse lung disorders^43,44^. Furthermore, systemic intravenous transplantation may be a suitable administration route in other future clinical scenarios. In our present study, intravenous IMRCs improved the survival rate of mice with BLM-induced lung injury in a dose-dependent manner, by inhibiting both pulmonary inflammation and fibrosis. As demonstrated in vitro, IMRCs inhibited the production of pro-inflammatory cytokines and pro-fibrosis cytokines, such as TNF-α and TGF-β1, in lung tissue in vivo. Furthermore, lung function indicators such as PV, IC, Rrs, Crs, Ers, Rn, G, H and FVC were improved compared with the BLM group.

It should be noted that IMRCs were administered at 1 and 7 days after initiation of BLM-injury, but before fibrosis has fully developed. In fact several studies have reported that primary MSCs treatments do not improve pre-established pulmonary fibrosis, while some studies suggest that primary MSCs actually exacerbate pulmonary fibrosis^45^. Regarding the efficacy of IMRCs in established fibrosis, further studies are necessary and our own investigation is still ongoing. It is unclear whether primary MSCs do promote pulmonary fibrosis, but their lack of positive therapeutic effect may be due to differences in MSC subtypes and functions, fibrosis stages after induction, species of model animals, and routes of administration of MSCs. Thus, clinical applications with MSCs should be considered carefully. As noted in this study, it is important to carefully document the origin, preparation and characterization of MSC-like cells. Here we have established a substantial profile of efficacy and safety for the hESC-derived IMRCs. These efforts provide considerable grounds for hope that this artificial cell-type could provide significant clinical benefit in the treatment of inflammatory and fibrotic disease conditions.

## Conclusions

In summary, our findings show that intravenously delivered IMRCs inhibited both pulmonary inflammation and fibrosis after acute lung injury in vivo. IMRCs significantly improved the survival rate in a dose-dependent manner. Furthermore, the mechanism for amelioration of pulmonary injury may be mediated by the IMRCs’ paracrine action rather than their potential to differentiate and replace the damaged alveolar epithelial cells. The IMRCs’ functional inhibition of TGF-β1-induced fibrosis could also be part of its mechanism. IMRCs were superior to UCMSCs and pirfenidone in therapeutic efficacy against lung injury and fibrosis. Furthermore, IMRCs showed an excellent safety profile in both mice and monkeys. In light of recent public health crises involving pneumonia, respiratory failure, ALI and ARDS, our pre-clinical results indicate that IMRCs are ready to be carefully considered for human trials in the treatment of lung injury and fibrosis.

## Acknowledgements

This work was supported by Beijing Municipal Science & Technology Commission (Z181100003818005 to Qi Zhou), National Key Research and Development Program (2018YFA0108400 and 2017YFA0104403 to Jie Hao, 2017YFA0105001 and 2016YFA0101502 to Liu Wang, 2020YFC0843900 to Qi Zhou, 2020YFC0841900 to Baoyang Hu), the Strategic Priority Research Program of the Chinese Academy of Sciences (XDA16030701 to Liu Wang, XDA16030401 to Wei Li, XDA16040502 to Jie Hao), the National Natural Science Foundation of China (31621004 to Qi Zhou and Wei Li), the Key Research Projects of the Frontier Science of the Chinese Academy of Sciences (QYZDY-SSW-SMC002 to Qi Zhou), and China Postdoctoral Science Foundation (2019M660791 to Lei Wang).

## Author contributions

B.H., H.D. and J.H. conceived the project and supervised the experiments. These authors contributed equally: J.W., D.S., Z.L. and B.G.. J.W., D.S., Z.L. and B.G. wrote the manuscript with help from all the authors. J.W., D.S., Z.L., B.G., Y.X., W.L., L.L., C.F., T.G., Y.C., Y.L., Z.W., J.W., S.Y., P.L., L.W., Y.W., L.P., H.W., T.Z., G.N.S., Z.H., G.F., W.L., Y.H., R.J., N.SC., Q.Z., L.W., B.H., H.D. and J.H. participated in the experiments and data analysis.

## References

1. Yang, J. & Jia, Z. Cell-based therapy in lung regenerative medicine. Regenerative medicine research. 2, 7 (2014).

2. Fan, E., Brodie, D. & Slutsky, A. S. Acute Respiratory Distress Syndrome: Advances in Diagnosis and Treatment. Jama. 319, 698–710 (2018).

3. Rubenfeld, G. D. et al. Incidence and outcomes of acute lung injury. N Engl J Med. 353, 1685–1693 (2005).

4. Bellani, G. et al. Epidemiology, Patterns of Care, and Mortality for Patients With Acute Respiratory Distress Syndrome in Intensive Care Units in 50 Countries. Jama. 315, 788–800 (2016).

5. Hamacher, J. et al. Tumor necrosis factor-alpha and angiostatin are mediators of endothelial cytotoxicity in bronchoalveolar lavages of patients with acute respiratory distress syndrome. Am J Respir Crit Care Med. 166, 651–656 (2002).

6. Raghu, G. et al. An official ATS/ERS/JRS/ALAT statement: idiopathic pulmonary fibrosis: evidence-based guidelines for diagnosis and management. Am J Respir Crit Care Med. 183, 788–824 (2011).

7. Marshall, R., Bellingan, G. & Laurent, G. The acute respiratory distress syndrome: fibrosis in the fast lane. Thorax. 53, 815–817 (1998).

8. Dhainaut, J. F., Charpentier, J. & Chiche, J. D. Transforming growth factor-beta: a mediator of cell regulation in acute respiratory distress syndrome. Critical care medicine. 31, S258–264 (2003).

9. King, T. E., Jr. et al. A phase 3 trial of pirfenidone in patients with idiopathic pulmonary fibrosis. N Engl J Med. 370, 2083–2092 (2014).

10. Yoon, H. Y., Park, S., Kim, D. S. & Song, J. W. Efficacy and safety of nintedanib in advanced idiopathic pulmonary fibrosis. Respiratory research. 19, 203 (2018).

11. Zhou, H., Liu, M. & Lyu, X. [Research progress of mesenchymal stem cells in the treatment of ALI]. Zhonghua wei zhong bing ji jiu yi xue. 29, 1039–1042 (2017).

12. Serrano-Mollar, A. Cell Therapy in Idiopathic Pulmonary Fibrosis(†). Med Sci (Basel). 6, 64 (2018).

13. Ortiz, L. A. et al. Interleukin 1 receptor antagonist mediates the antiinflammatory and antifibrotic effect of mesenchymal stem cells during lung injury. Proceedings of the National Academy of Sciences of the United States of America. 104, 11002–11007 (2007).

14. Walter, J., Ware, L. B. & Matthay, M. A. Mesenchymal stem cells: mechanisms of potential therapeutic benefit in ARDS and sepsis. Lancet Respir Med. 2, 1016–1026 (2014).

15. G, W. J. et al. Mesenchymal stem (stromal) cells for treatment of ARDS: a phase 1 clinical trial. The Lancet. Respiratory medicine. 3 (2015).

16. Thomson, J. A. et al. Embryonic stem cell lines derived from human blastocysts. Science (New York, N.Y.). 282, 1145–1147 (1998).

17. Barberi, T., Willis, L. M., Socci, N. D. & Studer, L. Derivation of multipotent mesenchymal precursors from human embryonic stem cells. PLoS medicine. 2, e161 (2005).

18. Hwang, N. S. et al. In vivo commitment and functional tissue regeneration using human embryonic stem cell-derived mesenchymal cells. Proceedings of the National Academy of Sciences of the United States of America. 105, 20641–20646 (2008).

19. Vodyanik, M. A. et al. A mesoderm-derived precursor for mesenchymal stem and endothelial cells. Cell stem cell. 7, 718–729 (2010).

20. Gu, Q. et al. Accreditation of Biosafe Clinical-Grade Human Embryonic Stem Cells According to Chinese Regulations. Stem cell reports. 9, 366–380 (2017).

21. Pittenger, M. F. et al. Multilineage potential of adult human mesenchymal stem cells. Science (New York, N.Y.). 284, 143–147 (1999).

22. Pittenger, M. F. et al. Mesenchymal stem cell perspective: cell biology to clinical progress. NPJ Regenerative medicine. 4, 22 (2019).

23. Han, X. et al. Mapping human pluripotent stem cell differentiation pathways using high throughput single-cell RNA-sequencing. Genome biology. 19, 47 (2018).

24. Kim, D., Langmead, B. & Salzberg, S. L. HISAT: a fast spliced aligner with low memory requirements. Nature methods. 12, 357–360 (2015).

25. Trapnell, C. et al. Transcript assembly and quantification by RNA-Seq reveals unannotated transcripts and isoform switching during cell differentiation. Nature biotechnology. 28, 511–515 (2010).

26. Huang da, W., Sherman, B. T. & Lempicki, R. A. Systematic and integrative analysis of large gene lists using DAVID bioinformatics resources. Nature protocols. 4, 44–57 (2009).

27. Subramanian, A. et al. Gene set enrichment analysis: a knowledge-based approach for interpreting genome-wide expression profiles. Proceedings of the National Academy of Sciences of the United States of America. 102, 15545–15550 (2005).

28. Prasad, V. K. et al. Efficacy and safety of ex vivo cultured adult human mesenchymal stem cells (Prochymal) in pediatric patients with severe refractory acute graft-versus-host disease in a compassionate use study. Biology of blood and marrow transplantation : journal of the American Society for Blood and Marrow Transplantation. 17, 534–541 (2011).

29. Horibata, S., Vo, T. V., Subramanian, V., Thompson, P. R. & Coonrod, S. A. Utilization of the Soft Agar Colony Formation Assay to Identify Inhibitors of Tumorigenicity in Breast Cancer Cells. Journal of visualized experiments : JoVE, e52727 (2015).

30. Zhang, W. et al. SIRT6 deficiency results in developmental retardation in cynomolgus monkeys. Nature. 560, 661–665 (2018).

31. Quinlan, A. R. & Hall, I. M. BEDTools: a flexible suite of utilities for comparing genomic features. Bioinformatics (Oxford, England). 26, 841–842 (2010).

32. Berninger, M. T. et al. Fluorescence molecular tomography of DiR-labeled mesenchymal stem cell implants for osteochondral defect repair in rabbit knees. European radiology. 27, 1105–1113 (2017).

33. Yang, J. et al. Tumor tropism of intravenously injected human-induced pluripotent stem cell-derived neural stem cells and their gene therapy application in a metastatic breast cancer model. Stem cells (Dayton, Ohio). 30, 1021–1029 (2012).

34. Wagers, S., Lundblad, L., Moriya, H. T., Bates, J. H. & Irvin, C. G. Nonlinearity of respiratory mechanics during bronchoconstriction in mice with airway inflammation. Journal of applied physiology (Bethesda, Md. : 1985). 92, 1802–1807 (2002).

35. Overmyer, K. A., Thonusin, C., Qi, N. R., Burant, C. F. & Evans, C. R. Impact of anesthesia and euthanasia on metabolomics of mammalian tissues: studies in a C57BL/6J mouse model. PloS one. 10, e0117232 (2015).

36. Lian, Q. et al. Derivation of clinically compliant MSCs from CD105+, CD24-differentiated human ESCs. Stem cells (Dayton, Ohio). 25, 425–436 (2007).

37. Lee, E. J. et al. Novel embryoid body-based method to derive mesenchymal stem cells from human embryonic stem cells. Tissue engineering. Part A. 16, 705–715 (2010).

38. Shi, Y. et al. How mesenchymal stem cells interact with tissue immune responses. Trends in immunology. 33, 136–143 (2012).

39. Milani, S. et al. Differential expression of matrix-metalloproteinase-1 and -2 genes in normal and fibrotic human liver. The American journal of pathology. 144, 528–537 (1994).

40. Benyon, R. C., Iredale, J. P., Goddard, S., Winwood, P. J. & Arthur, M. J. Expression of tissue inhibitor of metalloproteinases 1 and 2 is increased in fibrotic human liver. Gastroenterology. 110, 821–831 (1996).

41. Du, C. et al. Transplantation of human matrix metalloproteinase-1 gene-modified bone marrow-derived mesenchymal stem cell attenuates CCL4-induced liver fibrosis in rats. International journal of molecular medicine. 41, 3175–3184 (2018).

42. Sgalla, G. et al. Idiopathic pulmonary fibrosis: pathogenesis and management. Respiratory research. 19, 32 (2018).

43. Zhu, Y. G., Hao, Q., Monsel, A., Feng, X. M. & Lee, J. W. Adult stem cells for acute lung injury: remaining questions and concerns. Respirology (Carlton, Vic.). 18, 744–756 (2013).

44. Kanelidis, A. J., Premer, C., Lopez, J., Balkan, W. & Hare, J. M. Route of Delivery Modulates the Efficacy of Mesenchymal Stem Cell Therapy for Myocardial Infarction: A Meta-Analysis of Preclinical Studies and Clinical Trials. Circulation research. 120, 1139–1150 (2017).

45. Weiss, D. J. & Ortiz, L. A. Cell therapy trials for lung diseases: progress and cautions. Am J Respir Crit Care Med. 188, 123–125 (2013).

