## supplemental information for "Immunity-and-Matrix-Regulatory Cells Derived from Human Embryonic Stem Cells Safely and Effectively Treat Mouse Lung Injury and Fibrosis"

**Supplementary Table S1 Biological safety analysis of the human embryonic stem cells (hESCs)-derived immunity- and matrix-regulatory cells (IMRCs)[a].**

| Sterility and pathogen[b] | IMRCs |
|---|---|
| **[Identification tests]** | |
| Cell morphology | Adherent cells in monolayer, showing fibroblast-like morphology |
| Isozyme analysis | B type of human origin |
| Short tandem repeats (STRs) | Expressing 16 STR loci, each STR locus has 1-2 alleles. STR data is consistent with its original hESCs. |
| **[Bacteria and fungi]** | Negative |
| **[Mycoplasma]** | Negative |
| **[Exogenous virus test - in vitro]** | |
| Cell observation | Cell morphology normal |
| Hemadsorption test | Negative |
| Hemagglutination test | Negative |
| **[Exogenous virus test - in vivo]** | |
| Cell inoculation in suckling mice | Survival rate > 80% |
| Cell inoculation in adult mice | Survival rate > 80% |
| Cell inoculation in guinea pigs | Survive, no tuberculosis |
| Cell inoculation in rabbits | Survive, no abnormality |
| Survival rate of 5- to 6-day-old chick embryos | Survival rate 100% |
| Survival rate of 9- to 11-day-old chick embryos | Survival rate 100% |
| Hemagglutination test of 9- to 11-day-old chick embryo allantoic fluid | Negative |
| **[Human virus test]** | |
| Human Immuno Deficiency Virus I (HIV-I) | Negative |
| Human Hepatitis B virus (HBV) | Negative |
| Human Hepatitis C virus (HCV) | Negative |

| | |
|---|---|
| Human Cytomegalo Virus (HCMV) | Negative |
| Epstein-Barr virus (EBV) | Negative |
| Human Papillomavirus (HPV) | Negative |
| Human herpes virus 6, 7 (HHV-6, 7) | Negative |
| Human Parvovirus B19 | Negative |
| **[Bovine virus test]** | |
| Bovine parvovirus | Negative |
| Bovine adenovirus | Negative |
| **[Porcine virus test]** | |
| Porcine parvovirus | Negative |
| Porcine torque teno virus | Negative |
| **[Retrovirus test]** | |
| Reverse transcriptase activity | Negative |
| **[Immunological response test]** | |
| Lymphocyte proliferation inhibition rate | 92.8%, coculture of IMRCs and peripheral blood monocytes (PBMCs) in the ratio of 1:5 |
| Specific subsets of lymphocytes test | |
| Th1 lymphocyte proliferation inhibition rate | 45.2%, coculture of IMRCs and PBMCs in the ratio of 1:5 |
| Th17 lymphocyte proliferation inhibition rate | 48.0%, coculture of IMRCs and PBMCs in the ratio of 1:5 |
| Treg lymphocyte proliferation inhibition rate | 13.8%, coculture of IMRCs and PBMCs in the ratio of 1:5 |
| TNF-α secretion of lymphocyte inhibition rate | 91.8%, coculture of IMRCs and PBMCs in the ratio of 1:5 |
| **[Biological effectiveness test]** | |
| CD73 (%) | 98.9 |
| CD90 (%) | 98.8 |
| CD105 (%) | 95.8 |

| | |
|---|---|
| CD29 (%) | 99.7 |
| HLA-ABC (%) | 99.5 |
| CD11 (%) | < 0.1 |
| CD19 (%) | < 0.1 |
| CD34 (%) | < 0.1 |
| CD45 (%) | < 0.1 |
| HLA-DR (%) | < 0.1 |
| **[Pluripotent cells residuals]** | |
| TRA-1-60$^{+}$ proportion assay by FACS | < 0.1% |
| SSEA-4$^{+}$ proportion assay by FACS | 0.3% |
| TRA-1-81$^{+}$ proportion assay by FACS | < 0.1% |
| OCT4 expression assay by immunofluorescence | Negative, while its original hESCs is positive |
| NANOG expression assay by immunofluorescence | Negative, while its original hESCs is positive |
| OCT4 expression assay by PCR | Negative, while its original hESCs is positive |
| Nanog expression assay by PCR | Negative, while its original hESCs is positive |
| Teratoma formation in SCID mice | Six weeks after cell inoculation in SCID mice, no teratoma formation was observed, while its original hESCs is positive |
| **[Biopreparate test]** | |
| Endotoxin assay | ≤ 0.5 EU/mL |
| Bovine serum albumin residuals | < 50 ng/mL |

[a]This table is translated from NIFDC report (NO. SH201905849).

[b]The "Pharmacopoeia of the People's Republic of China, Edition 2015, Volume III" was used as a reference for the testing methods.

### Figure S1

a

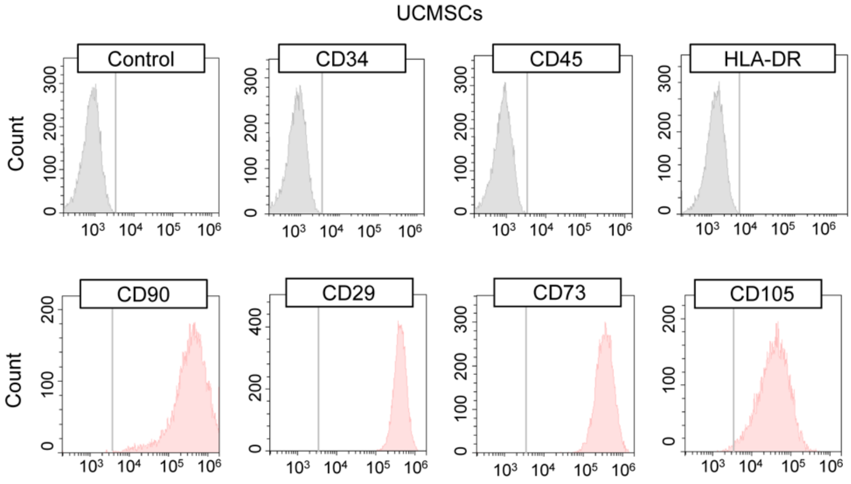


b

| | CD73 | CD90 | CD29 | CD105 | CD34 | CD45 | HLA-DR |
|---|---|---|---|---|---|---|---|
| IMRCs | 99.98±0.01 | 99.79±0.09 | 99.93±0.02 | 99.65±0.12 | 0.26±0.06 | 0.28±0.04 | 0.05±0.01 |
| UCMSCs | 99.76±0.12 | 96.46±0.32 | 99.97±0.01 | 97.73±0.06 | 0.03±0.01 | 0.44±0.15 | 0.02±0.01 |

c

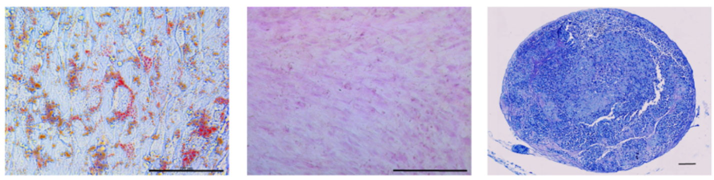


**Fig. S1 Derivation of Immunity- and Matrix-Regulatory Cells (IMRCs) from human embryonic stem cells (hESCs).**

**a** UCMSCs' expression of MSC-specific surface markers was determined by flow cytometry. Isotype control antibodies were used as controls for gating. The UCMSCs are $CD34^-/CD45^-/HLA\text{-}DR^-/CD90^+/CD29^+/CD73^+/CD105^+$ cells. **b** Quantification of flow cytometry for IMRCs and UCMSCs. **c** Representative staining of IMRCs after they were induced to undergo adipogenic differentiation (Oil Red O), osteogenic differentiation (Alizarin Red), and chondrogenic differentiation (Alcian Blue). Scale bars, 100 μm.

Figure S2

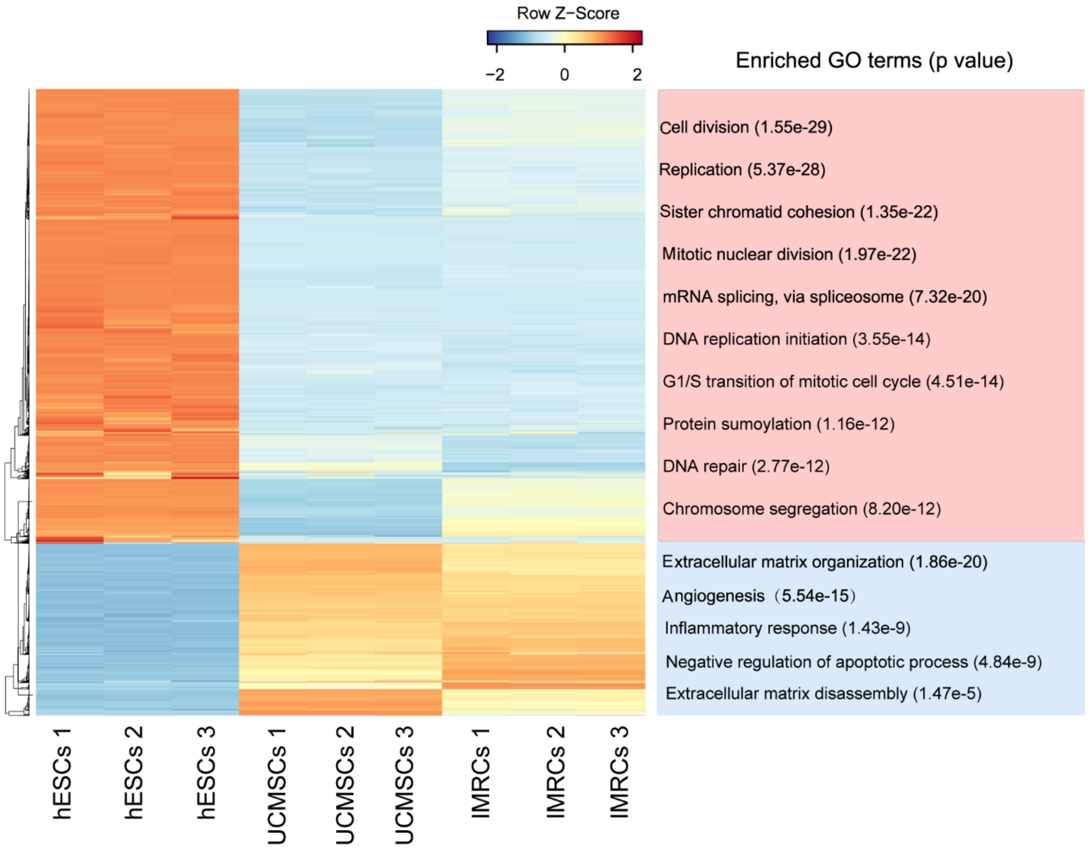


**Fig. S2 The differentially expressed genes among hESCs and UCMSCs, IMRCs.** In total, 4,730 differentially expressed genes were found in hESCs compared to UCMSCs and IMRCs. The enriched Gene Ontology (GO) terms and corresponding p values are shown.

Figure S3

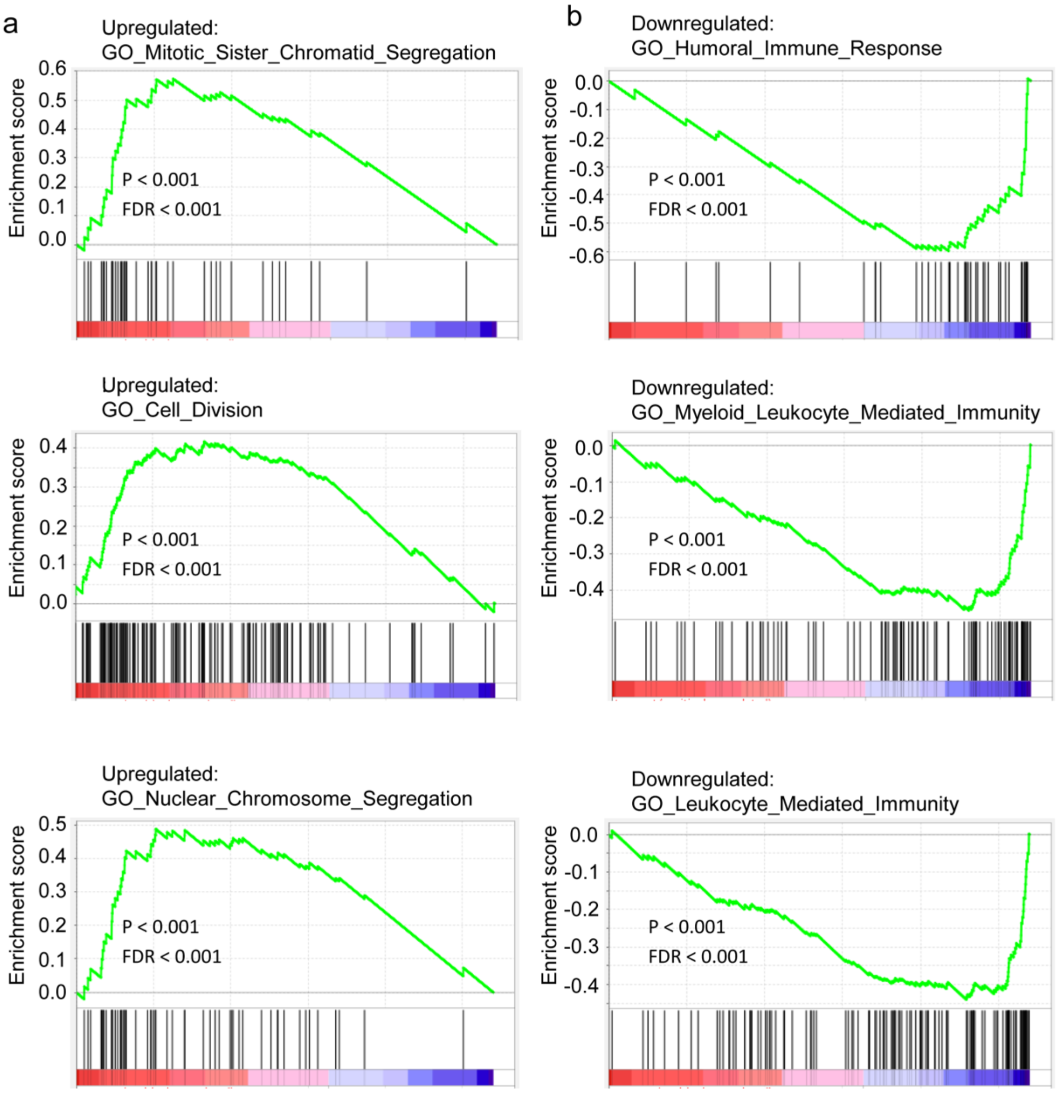


**Fig. S3 IMRCs possess unique gene expression characteristics.**

**a** Gene set enrichment analysis (GSEA) of the top up-regulated gene signature in IMRCs, compared with primary UCMSCs. **b** Gene set enrichment analysis (GSEA) of the top down-regulated gene signature in IMRCs, compared with primary UCMSCs.

Figure S4

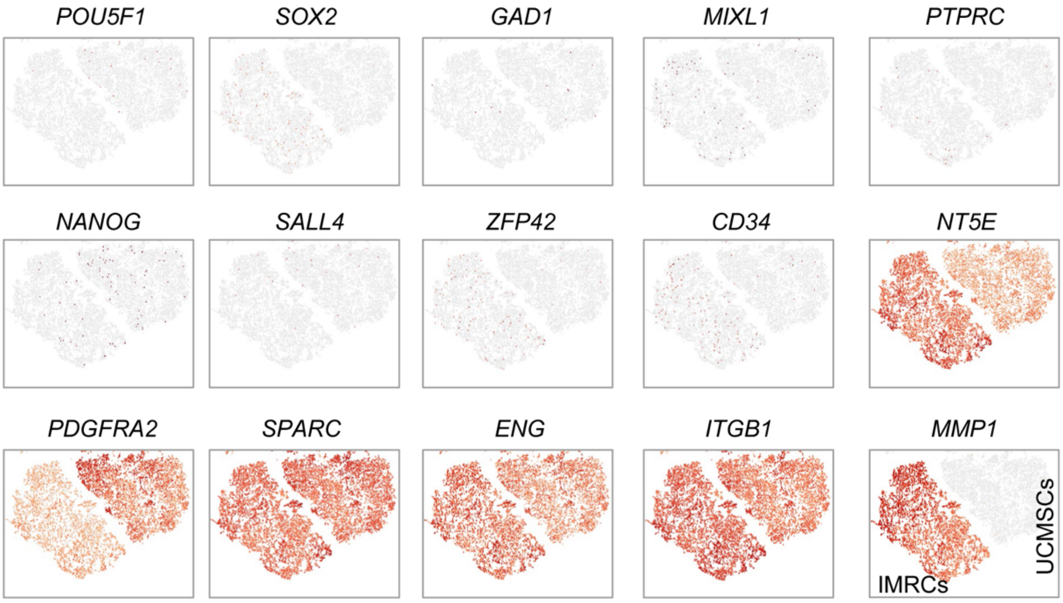


**Fig. S4 IMRCs possess unique gene expression characteristics.**

Heatmaps of specific gene expression amongst IMRCs and UCMSCs as measured by single cell RNAseq.

#### Figure S5

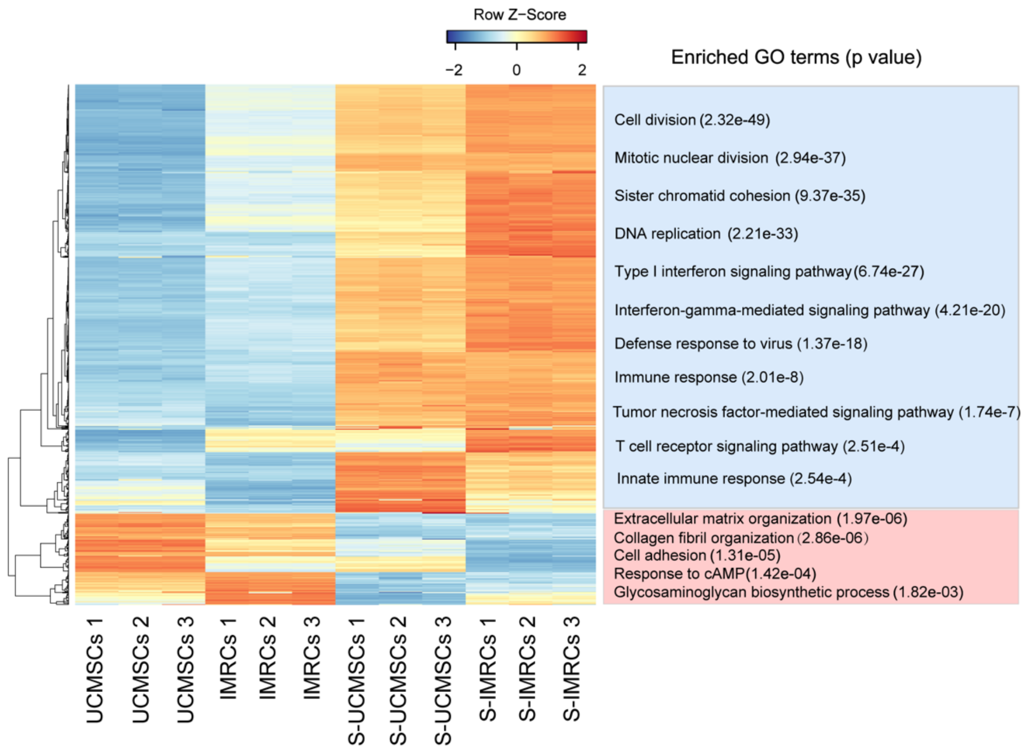


**Fig. S5 The differentially expressed genes in UCMSCs and IMRCs, before and after IFN-γ stimulation.** In total, 763 differentially expressed genes were found in UCMSCs and IMRCs before and after IFN-γ stimulation. The enriched Gene Ontology (GO) terms and corresponding p values are shown.

#### Figure S6

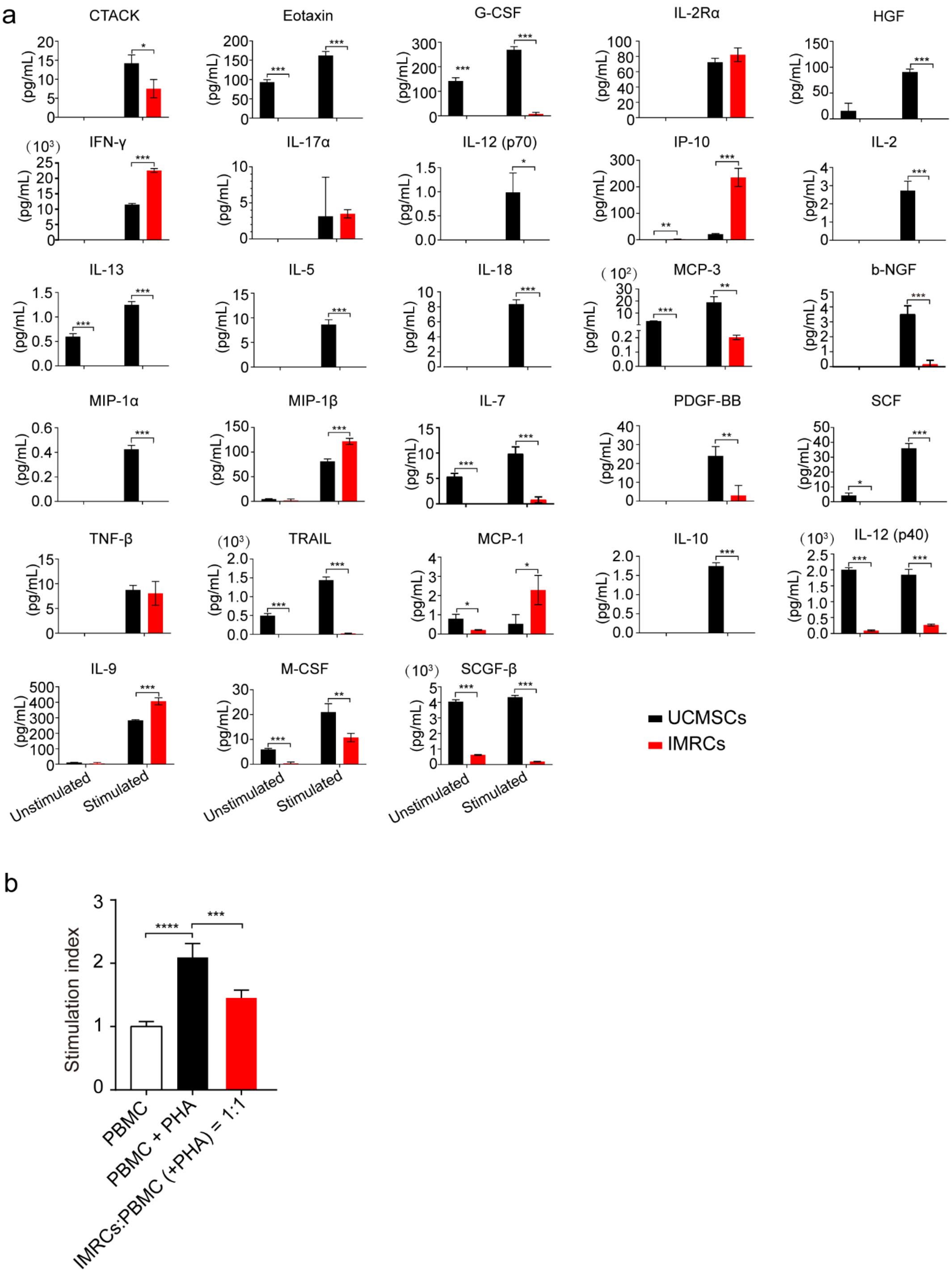


**Fig. S6 IMRCs activated by IFN-γ show hyper-immunomodulatory potency.**

**a** ELISA analysis of biologically relevant chemokines and cytokines in the secretomes of unstimulated or stimulated IMRCs and UCMSCs. bFGF, IL-15 and IL-16 were not detected. **b** The immunosuppressive effect of IMRCs on phytohaemagglutinin (PHA)-stimulated PBMCs' proliferation when cocultured together at a ratio of 1:1 ($2 \times 10^5$ IMRCs vs $2 \times 10^5$ PBMCs). * $p <$

0.05, ** $p < 0.01$, *** $p < 0.001$, **** $p < 0.0001$; data are represented as the mean ± SEM.

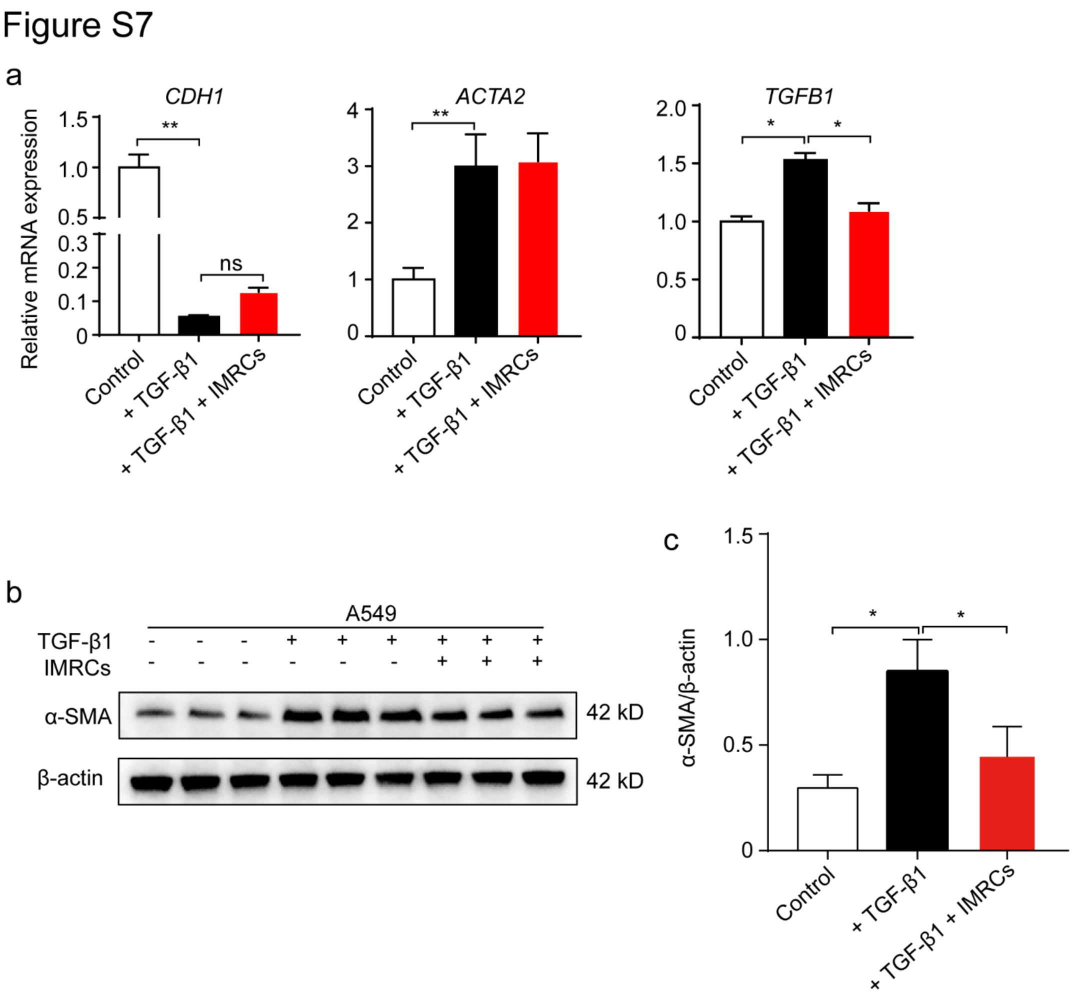


**Fig. S7 IMRCs reduce the pro-fibrotic effects of TGF-β1.**

**a** Quantitative PCR for *CDH1*, *ACAT2* and *TGFB1* mRNA in A549 cells, with or without 10 ng/mL TGF-β1 and IMRCs conditioned media treatment for 48 h. **b** Western blot for α-SMA protein expression in A549 cells, with or without 10 ng/mL TGF-β1 and IMRC conditioned media treatment for 48 h. **c** Quantification of the relative α-SMA protein expression levels in **(b)**. * $p < 0.05$, ** $p < 0.01$; data are represented as the mean ± SEM.

Figure S8

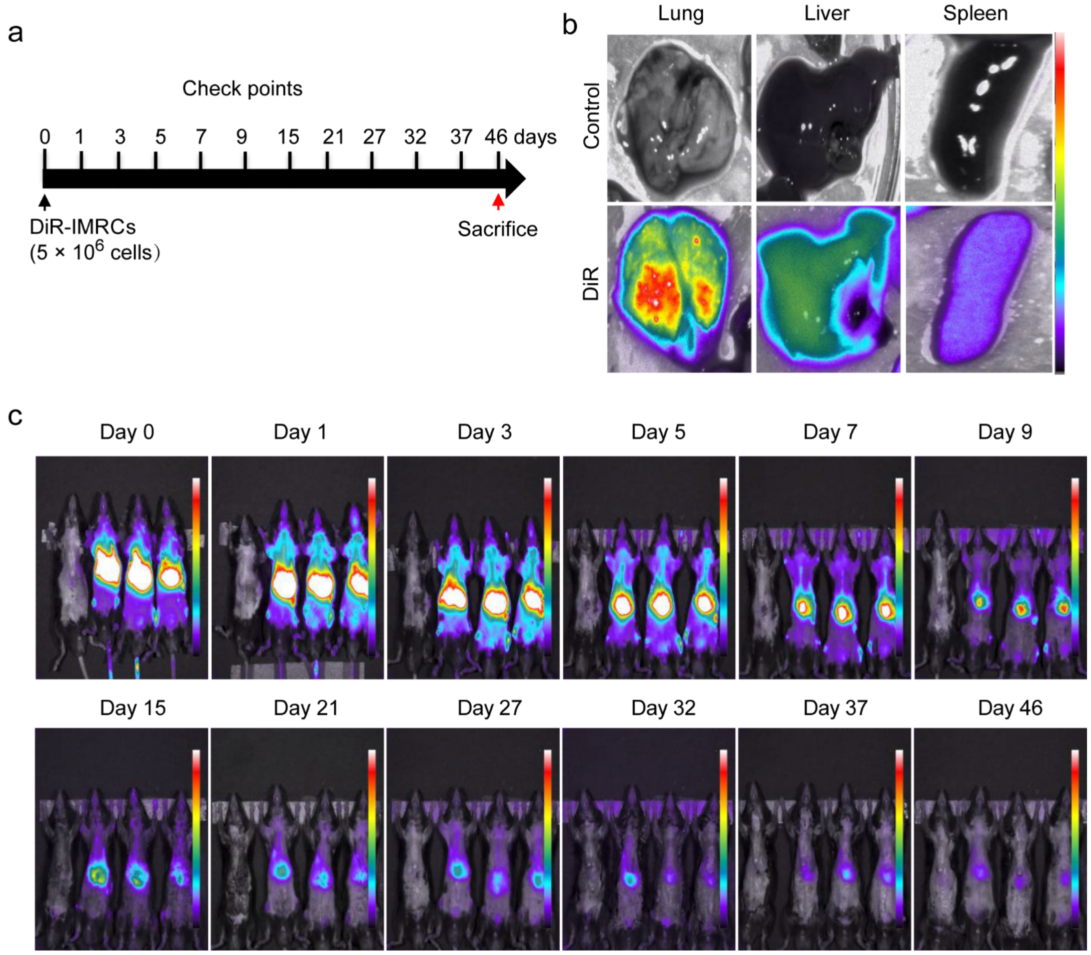


**Fig. S8 Evaluation of the safety of IMRCs transfusion.**

**a** Diagram of the animal experimental protocol. Mice were monitored using an in vivo imaging system at day 0, day 1, day 3, day 5, day 7, day 9, day 15, day 21, day 27, day 32, day 37 and day 46 after transplantation of DiR-labeled IMRCs. **b** In vivo imaging of the biodistribution of DiR far red fluorescence in the lung, liver and spleen, after injection of DiR-labeled IMRCs. **c** In vivo imaging of the biodistribution of DiR far red fluorescence after injection of DiR-labeled IMRCs.

#### Figure S9

a

Acute toxicity test in cynomolgus monkeys: safety analysis (after 6 months)

| Group | Number of monkey | Dose (cells/kg) | Weight | Ophthalmology examination | Vital signs |
|---|---|---|---|---|---|
| High dose | 1 | $1 \times 10^8$ | Normal | Normal | Normal |
| Medium dose | 1 | $0.26 \times 10^8$ | Normal | Normal | Normal |
| Low dose | 1 | $0.026 \times 10^8$ | Normal | Normal | Normal |

b

Long-term toxicity test in cynomolgus monkeys: safety analysis (one injection per week until 22 injections)

| Group | Number of monkey | Dose (cells/kg) | Weight | Body temperature | Food intake | All organs' weight |
|---|---|---|---|---|---|---|
| Saline | Female (3)<br>Male (3) | 10 mL / monkey | Normal | Normal | Normal | Normal |
| Low dose | Female (3)<br>Male (3) | $0.26 \times 10^7$ | Normal | Normal | Normal | Normal |
| High dose | Female (3)<br>Male (3) | $1 \times 10^8$ | Normal | Normal | Normal | Normal |

**Fig. S9 Evaluation of the safety of IMRCs transfusion.**

**a** Acute safety analysis of cynomolgus monkeys (*Macaca fascicularis*) injected with a low ($0.026 \times 10^8$), medium ($0.26 \times 10^8$) or high ($1 \times 10^8$) dose of IMRCs after 6 months. **b** Long-term safety analysis of cynomolgus monkeys injected with a low ($2.6 \times 10^6$) or high ($1 \times 10^8$) dose of IMRCs, or saline, once a week for 22 times.

Figure S10

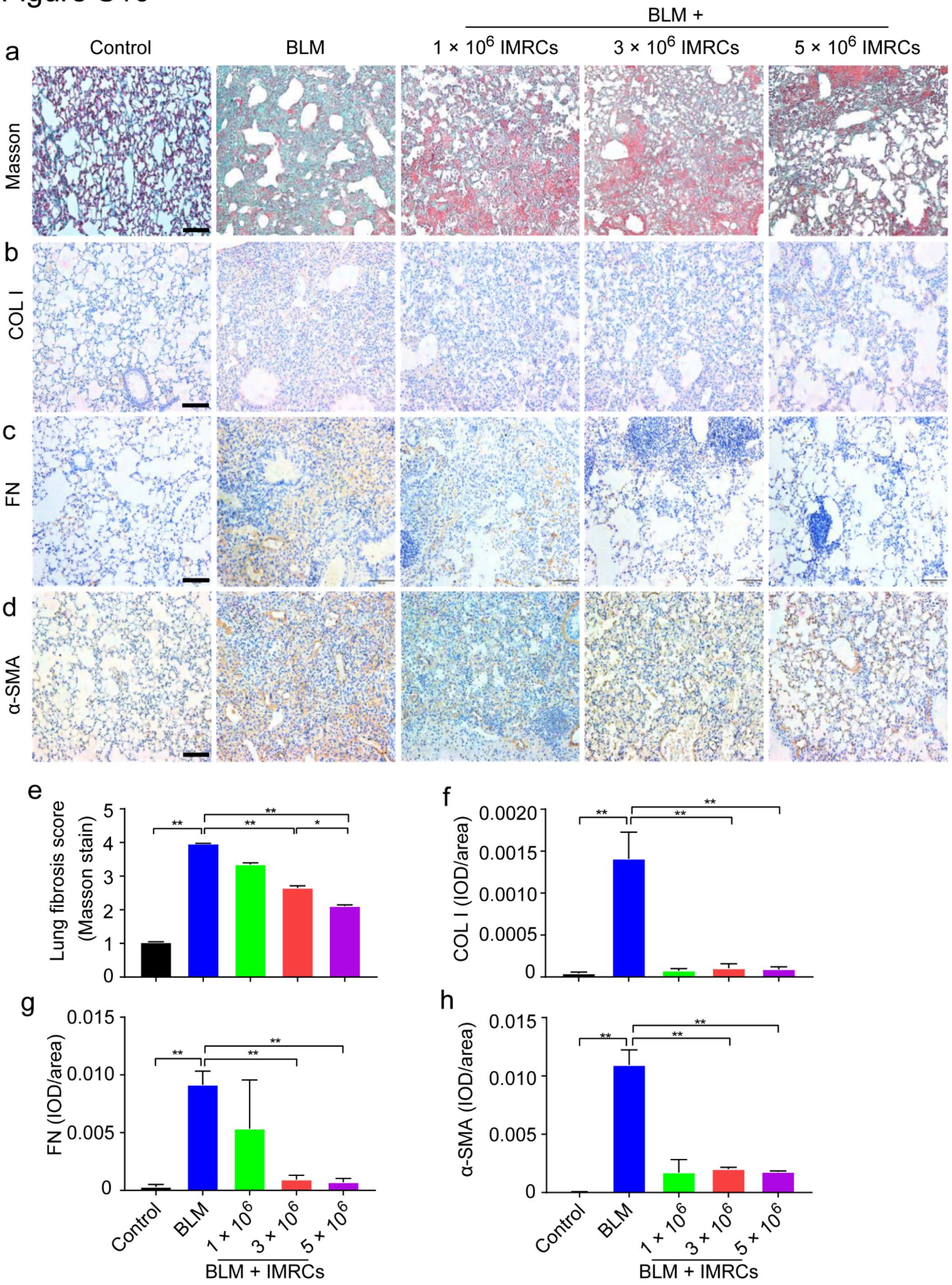


**Fig. S10 IMRC transfusion treats lung injury and fibrosis dose-dependently.**

**a** Representative lung sections of all treatment groups after Masson's trichrome staining (× 200). BLM, bleomycin. **b-d** Immunohistochemistry staining (brown) for the protein expression of **(b)** Collagen I (COL I), **(c)** Fibronectin (FN) and **(d)** α-smooth muscle actin (α-SMA), in mice receiving different interventions. **e** Lung fibrosis score, based on Masson's trichrome staining of lung sections in mice receiving different interventions. **f-h** Quantification of the immunohistochemistry staining for **(f)** Collagen I, **(g)** Fibronectin, and **(h)** α-SMA. * $p < 0.05$, ** $p < 0.01$; data are represented as the mean ± SEM.

Figure S11

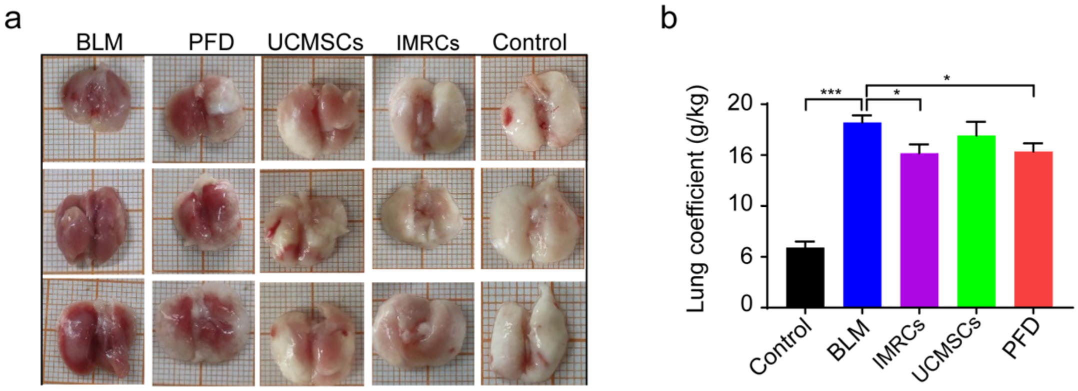


**Fig. S11 IMRC treatment of lung injury and fibrosis is superior to UCMSC and pirfenidone injections.**

**a** Representative images of whole lung from all treatment groups. One square, 1 mm. **b** Lung coefficient (wet lung weight/total body weight) of all treatment groups. * $p < 0.05$, ** $p < 0.01$, *** $p < 0.001$; data are represented as the mean ± SEM.

Figure S12

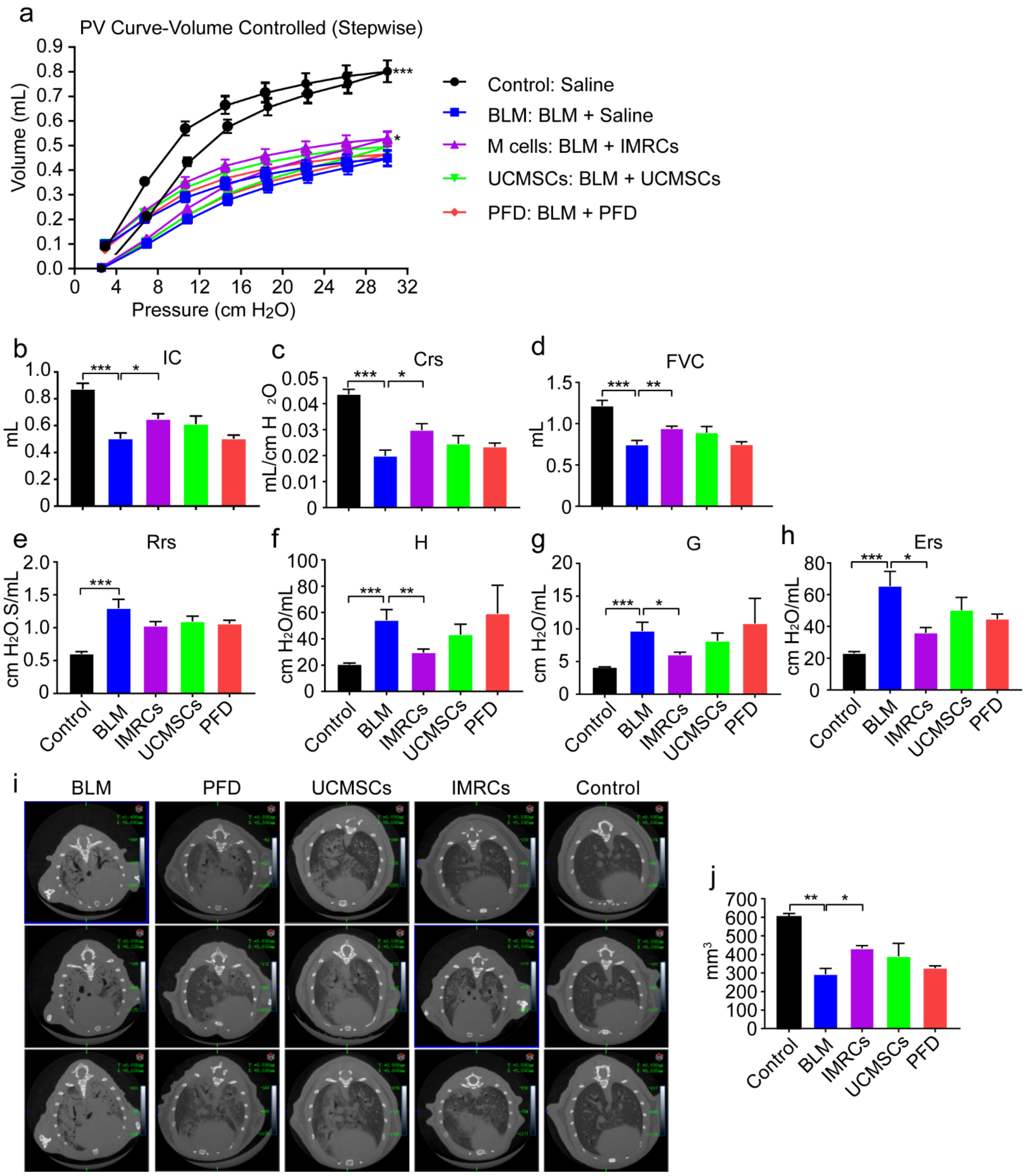


**Fig. S12 IMRC treatment of lung injury and fibrosis is superior to UCMSC and pirfenidone injections.**

**a-h** Lung mechanical function measured by FlexiVent at day 21, showing PV curves, IC, Rrs, Crs, Ers, G, H and FVC of all the experimental groups. PFD, pirfenidone; PV, pressure-volume; IC, inspiratory capacity; Rrs, respiratory resistance; Crs, static compliance; Ers, elastic resistance; G, tissue damping; H, tissue elasticity; FVC, forced vital capacity. **i-j** Micro-CT scans of the mouse lung and lung volume, in all the experimental groups. * $P < 0.05$; ** $P < 0.01$; *** $P < 0.001$.

**Supplementary Table S2** Primer sequences used in this study.

| Gene name | Forward (5'-3') | Reverse (5'-3') |
|---|---|---|
| *MMP1* | TGCTTCCCTGAGACCCAGTT | GATCACTTCTTTCCTTTGCATCAAG |
| *IDO1* | GCCAGCTTCGAGAAAGAGTTG | ATCCCAGAACTAGACGTGCAA |
| *CDH1* | AAAGGCCCATTTCCTAAAAACCT | TGCGTTCTCTATCCAGAGGCT |
| *ACTA2* | AAAAGACAGCTACGTGGGTGA | GCCATGTTCTATCGGGTACTTC |
| *COLLAGEN I* | GAGGGCCAAGACGAAGACATC | CAGATCACGTCATCGCACAAC |
| *COLLAGEN II* | TGGACGCCATGAAGGTTTTCT | TGGGAGCCAGATTGTCATCTC |
| *TGFβ1* | CTAATGGTGGAAACCCACAACG | TATCGCCAGGAATTGTTGCTG |
| *FIBRONECTIN* | AGGAAGCCGAGGTTTTAACTG | AGGACGCTCATAAGTGTCACC |
| *GAPDH* | CTCTGCTCCTCCTGTTCGAC | CGACCAAATCCGTTGACTCC |
